# Constraints on Persistent Activity in a Biologically Detailed Network Model of the Prefrontal Cortex with Heterogeneities

**DOI:** 10.1101/645663

**Authors:** Joachim Hass, Salva Ardid, Jason Sherfey, Nancy Kopell

## Abstract

Persistent activity, the maintenance of neural activation over short periods of time in cortical networks, is widely thought to underlie the cognitive function of working memory. A large body of modeling studies has reproduced this kind of activity using cell assemblies with strengthened synaptic connections. However, almost all of these studies have considered persistent activity within networks with homogeneous neurons and synapses, making it difficult to judge the validity of such model results for cortical dynamics, which is based on highly heterogeneous neurons. Here, we consider persistent activity in a detailed, strongly data-driven network model of the prefrontal cortex with heterogeneous neuron and synapse parameters. Surprisingly, persistent activity could not be reproduced in this model without incorporating further constraints. We identified three factors that prevent successful persistent activity: heterogeneity in the cell parameters of interneurons, heterogeneity in the parameters of short-term synaptic plasticity and heterogeneity in the synaptic weights. Our model predicts that persistent activity is recovered if the heterogeneity in the activity of individual interneurons is diminished, which could be achieved by a homeostatic plasticity mechanism. Such a plasticity scheme could also compensate the heterogeneities in the synaptic weights and short-term plasticity when applied to the inhibitory synapses. Cell assemblies shaped in this way may be potentially targeted by distinct inputs or become more responsive to specific tuning or spectral properties. Furthermore, the model predicts that a network that exhibits persistent activity is not able to dynamically produce intrinsic in vivo-like irregular activity at the same time, because heterogeneous synaptic connections are required for these dynamics. Thus, the background noise in such a network must either be produced by external input or constitutes an entirely different state of the network, which is brought about, e.g., by neuromodulation.

**Author summary:** To operate effectively in a constantly changing world, it is crucial to keep relevant information in mind for short periods of time. This ability, called working memory, is commonly assumed to rest on reverberating activity among members of cell assemblies. While effective in reproducing key results of working memory, most cell assembly models rest on major simplifications such as using the same parameters for all neurons and synapses, i.e., assuming homogeneity in these parameters. Here, we show that this homogeneity assumption is necessary for persistent activity to arise, specifically for inhibitory interneurons and synapses. Using a strongly data-driven network model of the prefrontal cortex, we show that the heterogeneities in the above parameters that are implied by *in vitro* studies prevent persistent activity. When homogeneity is imposed on inhibitory neurons and synapses, persistent activity is recovered. We propose that the homogeneity constraints can be implemented in the brain by means of homeostatic plasticity, a form of learning that keeps the activity of a network in a constant, homeostatic state. The model makes a number of predictions for biological networks, including a structural separation of networks responsible for generating persistent activity and spontaneous, noise-like activity.

## Introduction

Working memory can be defined as the ability to maintain a representation of a sensory stimulus, a thought or a memory retrieved from long-term storage over a period of time, even after the source of this representation is gone [1]. Working memory is a crucial prerequisite for many cognitive functions such as planning, reasoning, goal-directed behavior or language comprehension. It is also among the most commonly distorted cognitive functions in neurological and psychiatric disorders such as schizophrenia [2, 3]. A neural basis of working memory is widely believed to be persistent activity in cortical neurons, i.e., elevated firing rates that emerge during synaptic stimulation that persist after the input is removed [4–6]. This type of activity has been consistently found during working memory tasks in electrophysiological recordings [4, 7–9] as well as imaging recordings from humans [5, 10], most prominently in the prefrontal cortex (PFC). Furthermore, the stability of persistent activity is often strongly correlated with behavioral performance, i.e., if activity does not persist, animals are much more likely to make an error [4, 7].

The prominent role of persistent activity in working memory and cognition has inspired a large number of modeling studies that aim to understand the neural mechanisms underlying persistent activity [4, 6, 11]. The most widely studied of these mechanisms is reverberating synaptic activity in so-called attractor networks or cell assemblies, sets of neurons that are more strongly interconnected to each other compared to the average connectivity. This mechanism has been implemented in models of different levels of biological abstraction [4, 11–14]. In this study, we focus on attractor network models that rely on anatomically and physiologically stable assemblies, which may either be shaped by Hebbian learning or by proximity in space (either physical or defined by stimulus features), as opposed to more recent proposals of transient activations of cellular subsets [11], e.g., based on short-term synaptic modifications [15].

An important simplification in most of the existing models of persistent activity is disregard of the substantial heterogeneity of functional properties of neurons and synapses observed in the living brain, i.e., using the same set of parameters for all the simulated entities of the same type. Few studies have investigated the impact of incorporating heterogeneities into working memory models. The existing ones have focused on the ring model [16], for spatial working memory and heterogeneities in the parameters of pyramidal cells and their synaptic connections. Results suggest that random heterogeneities in the model cause the representation to drift to other spatial locations [17–19], while spatially structured heterogeneities [20], homeostatic regulation of pyramidal cell activity [17] and short-term synaptic plasticity [18, 19] can prevent this drift.

In this work, we investigate persistent activity of single cell assemblies within a biophysically detailed spiking network model of the PFC that is strongly constrained by in *vitro* data, including cellular and synaptic heterogeneities [21]. We find that these naturally occurring heterogeneities abolish bistability that is necessary for stimulus-dependent persistent activity. Thus, persistent activity itself is lost, instead of drifting away from the encoded location as in the ring model. Three aspects of heterogeneity are found to be harmful for persistent activity: Heterogeneous excitability of the interneurons (but not of the pyramidal cells, as in previous studies), heavy tails in the distribution of synaptic inputs [22] and heterogeneous short-term plasticity, i.e. different types of plasticity (facilitating, depressing or both) in synapses between neurons of the same type. We also show that persistent activity is recovered when the heterogeneities in the interneurons are compensated, e.g., by homeostatic plasticity [23, 24]. The same plasticity scheme could also compensate the heterogeneities in the synaptic weights and short-term plasticity when applied to the inhibitory synapses. However, as an important consequence of the homogeneity requirement for persistent activity, we show that cell assembly neurons cannot dynamically produce the often observed irregular ground state on their own: The dynamics that stabilize persistent activity are mutually exclusive with those dynamics necessary for the generation of irregular behavior. Instead, the assemblies may inherit these noise-like properties from input from other subnetworks or change its dynamic mode, e.g., using neuromodulation.

## Results

### No persistent activity in the detailed network model

Stimulus-dependent persistent activity is routinely seen in a wide variety of computational models that implement cell assemblies, sets of neurons that either innervate each other with a more dense connectivity or stronger synaptic weights. Mathematically, these assemblies show bistable behavior: In the absence of a stimulus, the firing rates of their member neurons are low, and there is no difference from the neurons outside the assembly (also called the spontaneous activity, e.g., [25]). When a stimulus is given to the neurons of the assembly, however, the strong activation triggers a positive feedback loop among the assembly neurons, which extends a high-rate persistent activity over time due to the long time constant of the NMDA currents. We attempted to generate persistent activity in the same way in a biologically validated model we recently proposed [21], which comprises broad, multivariate cell parameter distributions derived from *in vitro* recordings, laminar anatomy and experimentally constrained synaptic dynamics (see Methods for details). Surprisingly, we did not find the bistable behavior that is necessary for persistent activity as observed in vivo. Depending on the parameters of the model, either the spontaneous or the persistent activity state is stable, while the other one is not. Thus, the network either relaxes back to spontaneous activity after a transient reaction to the stimulus (Figure 1A) or persistent activity is present over the entire simulation time, only mildly modulated by the stimulus (Figure 1B). We varied a number of model parameters that are likely to affect persistent activity, such as the baseline synaptic weights between pyramidal cells and interneurons, synaptic weights and connectivity within the assembly (using scaling factors *s*_CA_ for weights and *p*_CA_ for connection probabilities), background currents into the pyramidal cells (*I*_ex_) and interneurons (*I*_inh_), as well as the size of the assembly *N*_CA_ and the ratio between the peak conductance of NMDA and AMPA currents, which is originally set to 1.09 [26, 27] (see Methods for a full overview over all parameter variations). Almost none of these manipulations led to persistent activity (with a single exception, which requires unphysiological levels of NMDA currents and puts the entire network in an epileptiform state with extremely high firing rates). Figure 1C shows the normalized activity in the cell assembly (see Methods for details) before the stimulus presentation (blue) and for the last 500 ms of the simulation (red). The results are sorted for ascending values of the activity before the stimulus. One can see that all possible activity values are covered, and the activity values at the end of the simulation hover around those before stimulation, i.e. activity that is low before stimulation stays low afterwards (Figure 1A), while high activity levels at the end of the simulation are already generated before stimulus presentation (Figure 1B), reflecting a continuum of monostable states.

**Fig 1.**
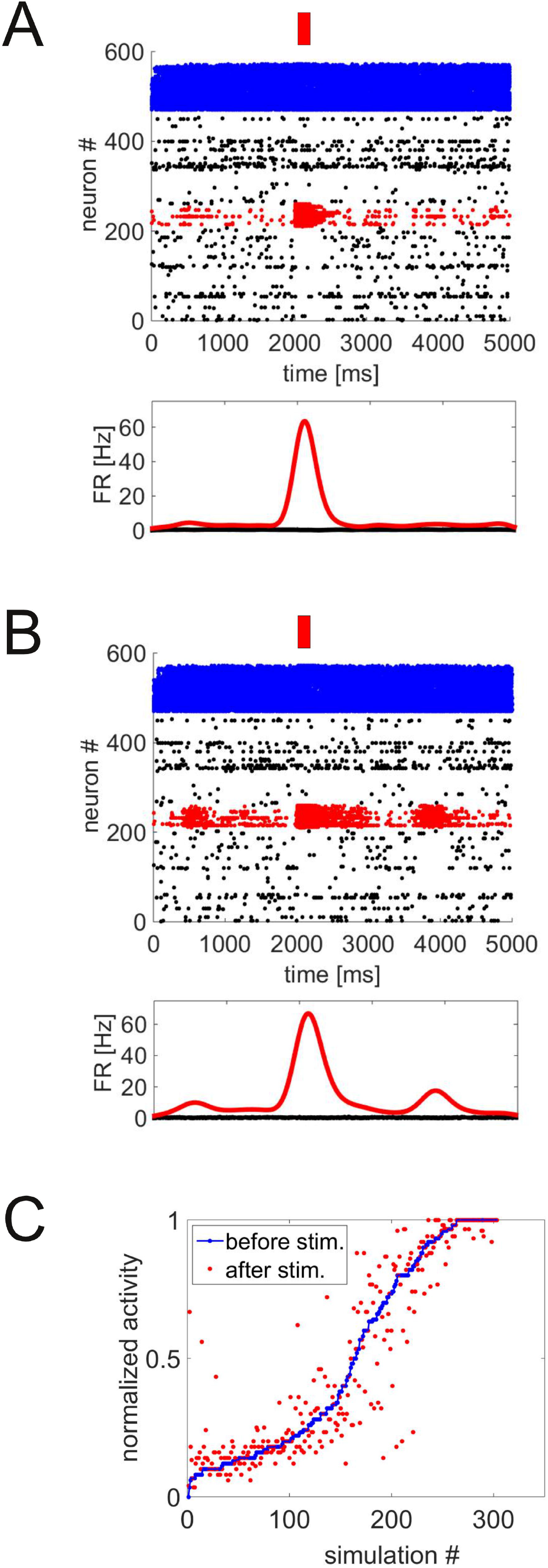
Failed persistent activity in the original network. A: Raster plot (upper panel) and instantaneous firing rate (lower panel) of layer 2/3 of the network, with pyramidal cells in red (cell assembly members) and black (neurons outside the assembly) and interneurons in blue. The red bar denotes a stimulus to the assembly neurons. Activity in the cell assembly quickly decays back to baseline after a few hundred ms. B: Raster plot as in A. Activity in the assembly persists, but arises spontaneously, i.e., not in reaction to the external stimulus. C: Summary of normalized activity in the cell assembly (see Methods for details) before stimulus presentation (blue) and at the end of the simulation (red) for 300 simulations with different parameters (assembly size between 30 and 80 cells). Simulations are sorted by ascending values of activity before the stimulus.

The remainder of the paper aims to identify how bistability can be introduced in a physiologically realistic network configuration. To differentiate bistability-induced persistent activity from the monostable states described above (Figure 1A and B), we introduce the measure *d*_PA_, which is defined as the difference between the normalized activity in the cell assembly after and before the stimulus, ranging between zero and one (*d*_PA_ is set to zero for negative differences, i.e. higher activity before the stimulus, see Methods for details). *d*_PA_ is close to zero in all of the monostable cases (largely constant firing rates over time) and increases only if the stimulus induces a persistent high-activity state, i.e., in the bistable case.

### Constraints for persistent activity

The above results (Figure 1) show that none of the simulations exploring a wide range of parameter combinations for the biologically validated network model was able to generate realistic persistent activity. This is a surprising result, as simpler models easily incorporated persistent activity in cell assemblies. Thus, we hypothesized that some of the biological complexity that is mimicked in the present model, but was missing in others, prevents persistent activity. To test this, we simplified the model to see which of the added features cause the problem. Most notably, we reduced the network to contain pyramidal cells and fast-spiking interneurons in layer 2/3 only (i.e., lacking layer 5 and four other types of interneurons, see Methods for details), used a uniform connection probability of 10%, removed short-term plasticity and reduced the input currents to the mean rheobase of each neuron type, which strongly reduced spontaneous firing. Furthermore, we made the neuron parameters of a given cell type much more uniform, reducing its standard deviation to 5% of the original value. These simplifications made the model more similar to previously studied models of persistent activity, in particular the Brunel and Wang model [25] (we also tested a configuration that completely mimicked that model, but results did not qualitatively change).

Most of these manipulations did not improve the ability of the network to produce persistent activity, with three exceptions: The reduction of the standard deviation of the neuron parameters, the removal of short-term plasticity and the use of a uniform connectivity across cell types. Each of these manipulations represents aspects of heterogeneity and variability of the network: Heterogeneity in the neuron parameters, the synaptic connectivity and the dynamics of synaptic response. In the following, we will investigate how these three aspects of heterogeneity relate to persistent activity and how they can be compensated in a biologically meaningful way.

#### Low-rheobase interneurons suppress persistent activity

First, we varied the amount of heterogeneity in the neuron parameters by jointly decreasing the standard deviations of the parameter distributions obtained from *in vitro* recordings to a given fraction of their original values for all parameters. Figure 2A shows the persistent activity measure *d*_PA_ as a function of cell parameter variability. Persistent activity emerges when variability is reduced to about 25% or less of the original value. This is indicated by values of *d*_PA_ above 0.3 (dotted line in Figure 2A), which has proven a good demarcation between successful and failed persistent activity using visual inspection of the raster plots, and will be used in this way throughout the paper. The exact choice of this demarcation is not critical for the results. To understand why the naturally occurring heterogeneity in the neuron parameters [21, 28] prevents persistent activity, we first analyzed the effect of individual cell types and neuron parameters, in particular the rheobase, the minimal amount of input current that elicits an action potential. Figure 2A shows the persistent activity measure *d*_PA_ as the cellular variability of pyramidal cells or interneurons were varied, respectively, while the other cells were left at their original parameter distribution. It is apparent that only the heterogeneity in the interneurons affects persistent activity (blue curve), while pyramidal cell variability does not have a noticeable effect, either when varied alone (red curve) or in concert with interneuron variability (black vs. blue curve).

**Fig 2.**
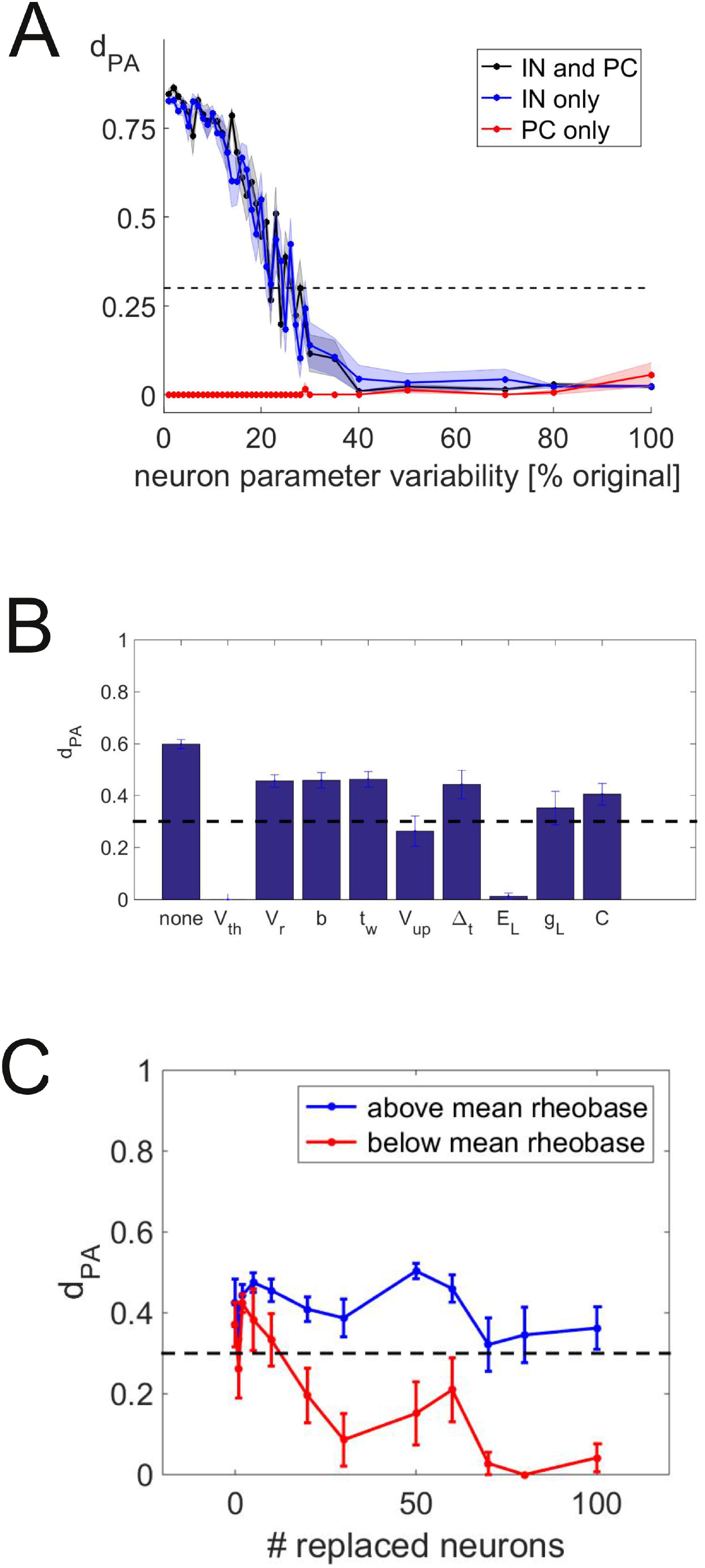
Breaking persistent activity by interneuron heterogeneity. A: *d*_PA_ as a function of the variability in neuron parameters in pyramidal cells only (red), interneurons only (blue) or all neurons (black). B: Persistent activity measure *d*_PA_ as individual parameters are increased back to 100% of their original variability, with all other parameters at 5%. The leftmost bar shows baseline *d*_PA_ if all parameters’ variability is left at 5%. The dashed line demarks the boundary between successful (*d*_PA_ ≥ 0.3) and failed (*d*_PA_ < 0.3) persistent activity. C: *d*_PA_ in simulations with interneuron variability set to 5% as increasing number of interneurons are replaced by neurons at the original variability, but only with rheobases above (blue) or below (red) the mean rheobase. Background currents are adjusted accordingly (see text for details). The dashed line demarks *d*_PA_ = 0.3 as in A.

To differentiate the effects of the individual neuron parameters of the interneurons, we started from a simulation where the variability of all parameters was set to 5% of their original values (Figure 2B, left bar, *d*_PA_ = 0.60 ± 0.02; mean ± SEM). Then, we replaced the reduced distribution of each of the nine independent model parameters of the neuron model (the simplified exponential integrate and fire neuron, or simpAdEx [28]) by the full distribution, one at a time. *d*_PA_ significantly decreased relative to baseline for each of these manipulations, but mostly stayed above 0.3, indicating successful persistent activity (dotted line in Figure 2B). However, *d*_PA_ dropped to values of almost zero when the original heterogeneity was used in the parameters *V*_th_ (onset of exponential upswing in the membrane potential, *d*_PA_ = 0.0 for all simulations) and *E_L_* (reversal potential of the leak current, *d*_PA_ = 0.01 ± 0.01; mean ± SEM).

The parameters *V*_th_ and *E_L_* that proved most important in the above analysis are the major determinants of the rheobase current in the simpAdEx [28]. Thus, we hypothesized that interneurons with low rheobase and high spontaneous firing rate will prevent the initiation of persistent activity. To test this hypothesis, we set up a simulation with two populations of interneurons, one using the original distribution and another with the variability of all parameters reduced to 5%. We first drew all interneurons from the reduced distribution and then replaced increasing numbers of interneurons by those drawn from the original distribution and computed the *d*_PA_ measure in each of these cases. Importantly, we replaced only neurons with rheobases above the average in one set of simulations (blue curve in Figure 2C) and only those with rheobases below the average in another one (red curve in Figure 2C). Background currents were adjusted such that the input relative to the mean rheobase was kept constant as neurons were replaced, so the effective mean input into the interneurons are not changed. While the replacement of neurons above rheobase does not affect persistent activity, *d*_PA_ drops as the number of replaced neurons below rheobase is increased. This implies that interneurons with rheobases below the population average dominate inhibition and prevent the initiation of persistent activity. Importantly, the adjustment of the (constant) background current was not sufficient to compensate the increased inhibitory activity. This suggests that the low-rheobase interneurons may induce a skew in the distribution of inhibitory activity rather than just shifting its mean.

#### Homeostatic compensation for rheobase heterogeneity recovers persistent activity

Persistent activity occurs in prefrontal cortex where cells are highly heterogeneous, so it is important to explain how persistent activity and cellular heterogeneity can be reconciled in a model of the prefrontal cortex. As we identified high spontaneous firing rates in low-rheobase interneurons as the main limiting factor for the initiation of persistent activity, the effect of these interneurons on total inhibition needs to be limited in a biologically realistic way. A potential mechanism limiting the maximal inhibition is homeostatic scaling of synaptic weights that aims to keep roughly uniform firing rates. This kind of homeostatic plasticity has been observed in both *in vivo* and *in vitro* studies and has been shown to regulate both excitatory and inhibitory synapses [24]. We emulate this homeostatic regulation by adjusting the excitatory background inputs (which are meant to represent the average synaptic input from outside the simulated network) individually for each interneuron such that the effective drive (input current minus rheobase) is the same for each interneuron. External input constitutes the bulk of the excitatory input to each cell, so extending the homeostasis to all excitatory input does not significantly change the results. With this adjustment, the distribution of firing rates in the network peaks at the mean rheobase, while previously, it was strongly skewed towards low-rheobase interneurons at high rates (Figure 3A). Furthermore, persistent activity emerges as the background currents are slightly decreased compared to the original value of 37pA (blue curve in Figure 3B). Importantly, the same manipulation lacks any effect without rheobase compensation (red curve). Thus, homeostatic plasticity is a possible way to compensate rheobase heterogeneity and allow for persistent activity.

**Fig 3.**
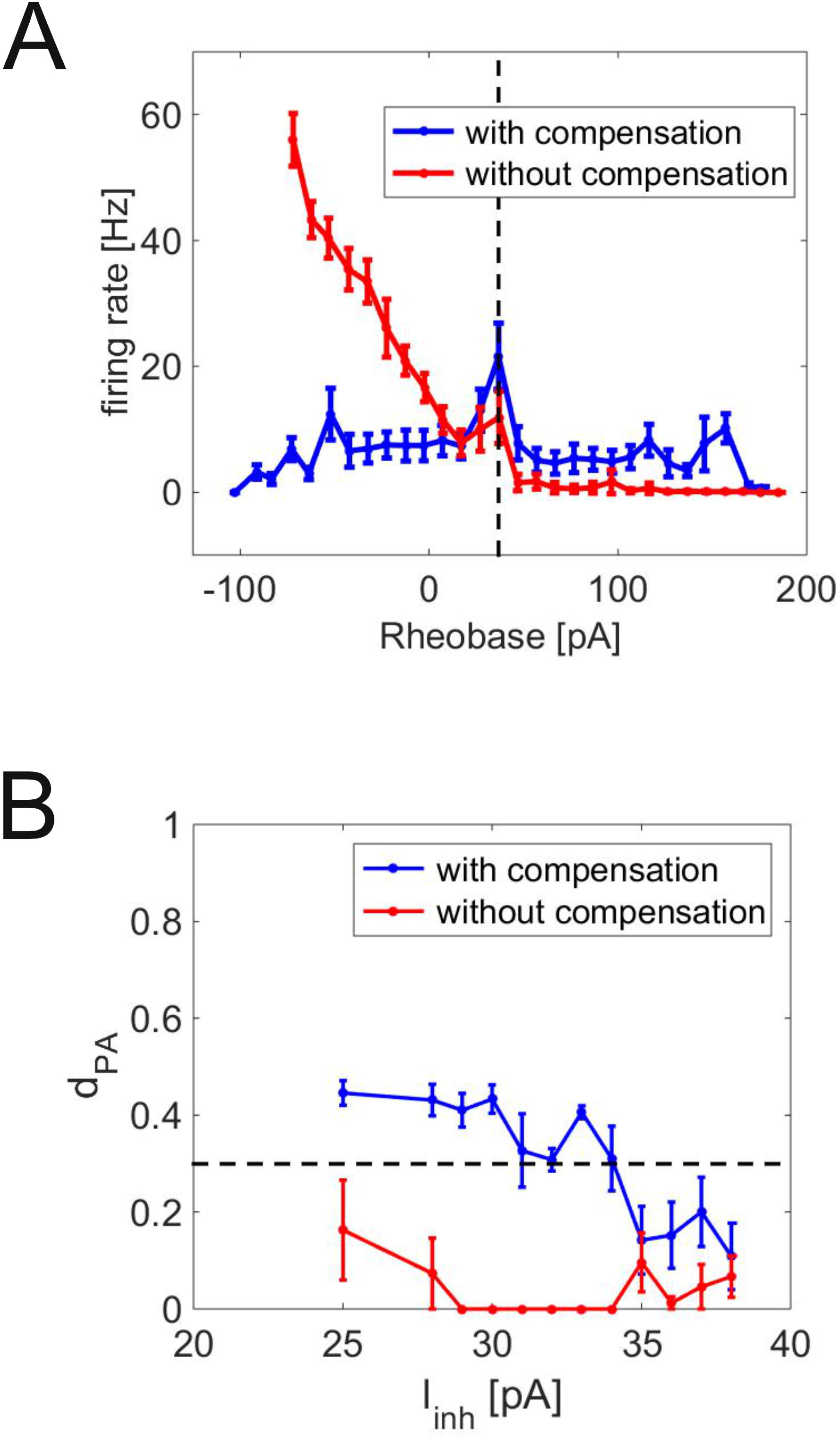
Compensation of rheobases restores persistent activity. A: Average interneuron firing rate as a function of rheobase with (blue) and without (red) background currents adapted to rheobases (mean ± SEM over all neurons). B: Persistent activity measure *d*_PA_ as a function of the background current into interneurons with (blue) and without (red) compensation of rheobases by adapting the background currents such that the input relative to each individual rheobase is constant (mean ± SEM over 12 simulations).

#### The role of short-term plasticity

The simulations shown in Figure 2 and 3 were conducted without short-term synaptic plasticity (STP). Here, we investigate the how the incorporation of STP affects persistent activity. STP refers to the dynamic change of synaptic efficacies in response to incoming spikes, which relaxes back to a baseline within several hundred milliseconds [29] (see Methods for details). This short time scale sets STP apart from long-term plasticity, which induces lasting changes in the synaptic weights. It has also been suggested that STP can be used as a dynamic basis for persistent activity [15], a proposal we do not consider in this study. Instead, we investigate the effect of STP heterogeneity on persistent activity generated by reverberation in a cell assembly. Our original model [21] contained three types of STP, which are suggested by electrophysiological data: Predominantly facilitating plasticity, predominantly depressing plasticity, and a mixed plasticity type [30, 31]. These studies also show that the cell types of the pre- and postsynaptic neuron determines the plasticity type of a given synapse. Connections among pyramidal cells, among fast-spiking interneurons and between fast-spiking interneurons and pyramidal cells, however, can exhibit all three types of plasticity with different probabilities (see Methods for details), introducing another factor of heterogeneity. We refer to simulations with this full distribution of STP types as “heterogeneous STP” and to those with only one STP type for a given pair of cell types as “homogeneous STP”. In the homogeneous case, only the most frequently occurring STP is being used for each pair of cell types (facilitating STP for pyramidal-pyramidal connections and depressing STP for fast-spiking interneurons inhibiting pyramidal cells as well as for interneuron-interneuron connections, c.f. Figure 2B in [21]).

The different curves in Figure 4 represent these different parameter configurations of STP: Heterogeneous STP (red curve), no STP at all (black curve) and homogeneous STP (blue curve). With heterogeneous STP (red curve), *d*_PA_ is close to zero for all values of cell parameter variability, while in the simulations without STP (black curve), persistent activity emerges as cellular heterogeneity is reduced. However, removing STP is not a necessary condition for persistent activity. Homogeneous STP (blue curve) also leads to persistent activity for low cell parameter heterogeneity. Thus, it is the heterogeneity of STP that prevents persistent activity to emerge, not STP per se.

**Fig 4.**
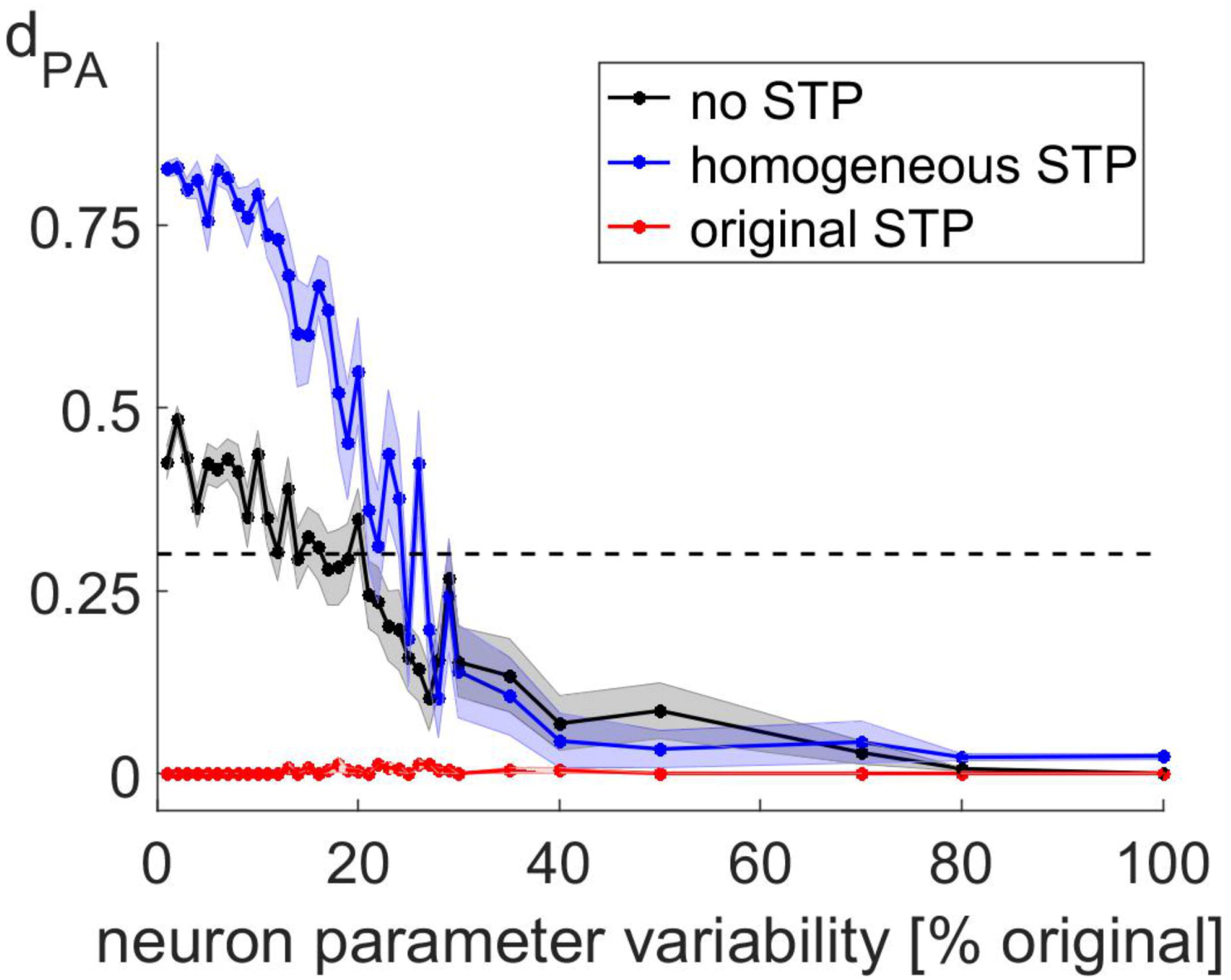
Interaction of cellular heterogeneity and short-term plasticity. Persistent activity measure *d*_PA_ as a function of the variability in neuron parameters with original STP (red), homogeneous STP (blue) and no STP (black). Mean *d*_PA_± SEM over 24 simulations is shown.

#### Dynamic impact of heterogeneous synaptic input

The above results show that persistent activity can be achieved when heterogeneities in interneuron cell parameters and short-term plasticity are removed or compensated. While these conditions are necessary for persistent activity, they are not sufficient: All of the simulations so far used a uniform connectivity of 10% across all cell types, while persistent activity could not be generated in the full network with all layers and cell types included and the original connectivity pattern obtained from in *vitro* data [21]. In this original pattern, connection probabilities are higher among interneurons (25%) and between interneurons and pyramidal cells (15% to 70%) compared to the connectivity among pyramidal cells (about 10%). Fast-spiking interneurons and Martinotti cells project particularly frequently onto pyramidal cells (about 50% and 70%, respectively). In the following, we investigate how this pattern of heterogeneous connectivities between different cell types shapes the dynamics of the full network (including homogeneous short-term plasticity) and why it prevents persistent activity from emerging.

First, we found that the most notable difference between the heterogeneous connectivity patterns and the homogeneous one (10% connectivity among all cell types) lies in the distribution of synaptic weights: In the heterogeneous connection pattern, pyramidal cells receive many more connections from fast-spiking interneurons and Martinotti cells in layer 2/3 compared to other cells types and cells from layer 5. Furthermore, synaptic weights for these densely connected cells type combinations are much stronger compared to others (2.30 mV and 1.91 mV, for fast-spiking interneurons and Martinotti cells, respectively, compared to an average of 0.10 mV for all other types of interneurons). These two sources of inhibition result in a bimodal distribution of synaptic weights (Figure 5A, red curve). When all connectivities are set to a uniform 10%, far fewer of the strong weights from fast-spiking interneurons and Martinotti cells are being drawn, while the distribution of the weaker weights is much less affected (Figure 5A, blue curve). This leads to greatly decreased numbers of strong inhibitory synapses onto L2/3 pyramidal cells in the uniform network compared to the heterogeneous one.

**Fig 5.**
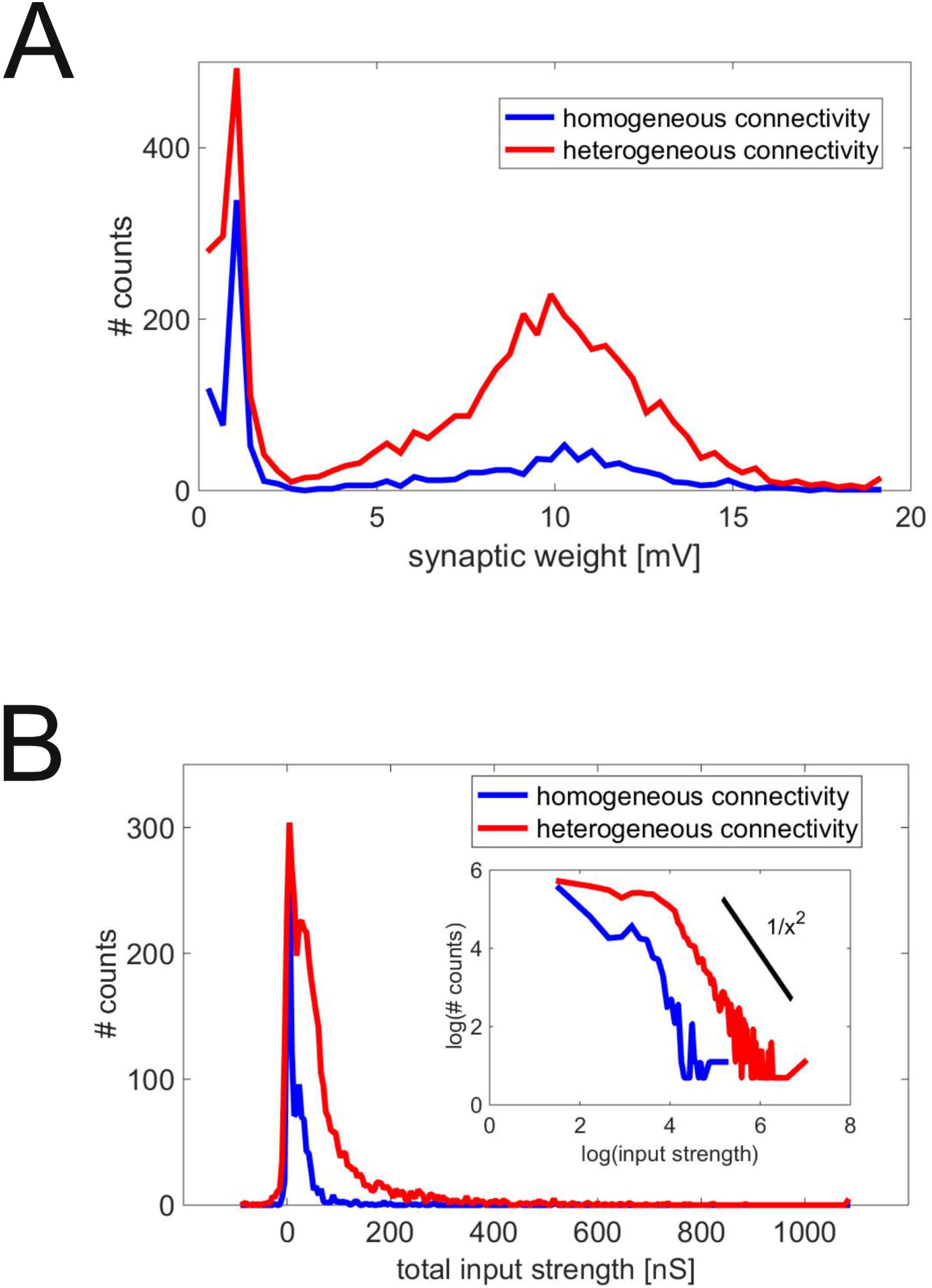
Dynamic difference between homogeneous and heterogeneous synaptic connections. A: Histogram of synaptic weights in networks with homogeneous connectivity (10% for all cell types, blue curve) and heterogeneous connectivity (connectivity constrained by data, red curve). B: Histogram of the total strength of each synapse in the networks with homogeneous and heterogeneous connectivity, computed as the product of the synaptic weight, the variables of short-term synaptic plasticity, averaged over all synaptic events, and the number of synaptic events. The inset shows the distributions on a log-log scale to demonstrate the heavy tail of the distributions. The black bar shows a 1/*x*^2^ distribution for comparison.

Second, we investigated the impact of the weight distribution on the total synaptic input from a given interneuron to a given cell assembly member. The total synaptic input is defined as the sum of synaptic conductance changes in the target neuron by spikes from the presynaptic interneuron, summed over the entire spike train of that interneuron. These total inputs follow of a heavy-tailed distribution (Figure 5B), where the tail sits on top of barely visible bimodal distribution at lower values. While the two modes reflect the distribution of the weights, the heavy tail occurs from the fact that large values of the total input require the rare coincidence of both large weights and high firing rates. Furthermore, inputs from interneurons with high rates are often strongly attenuated by short-term synaptic plasticity, as the predominant plasticity type from interneurons to pyramidal cells is synaptic depression. Thus, a high total input also requires suitable parameters of short-term plasticity, most importantly, a short time constant of synaptic depression. All these conditions only rarely coincide, but when they do, very large total synaptic inputs result, constituting the heavy tail of the distribution. Importantly, coincidence is much more likely if there is a large fraction of strong synapses, as in the noise network. Thus, the tail is heavier for heterogeneous connectivity (Figure 5B, red curve) compared to uniform connectivity (Figure 5B, blue curve).

Cell assembly neurons that receive even a few of those extreme inputs well show inhibitory conductances which are much higher than those from neurons which inhibitory inputs are all close to the mean, being effectively dominated by those few inputs. Thus, because of the much higher probability of extreme inputs under heterogeneous connectivity, we expected to see an increase both in the mean and in the standard deviations of the distribution of inhibitory conductances compared to uniform connectivity. Indeed, inhibitory conductances did show a higher mean and standard deviation across assembly neurons for the heterogeneous connectivity (54.30 ± 12.66 for heterogeneous connectivity compared to 14.25 ± 7.91 for uniform connectivity; mean ± SD; p=1.4 · 10^−54^, t(158)=24.01, two-sided t-test), while excitatory conductances only slightly (but significantly) differ in both configurations (1.05 ± 0.41 for heterogeneous connectivity compared to 1.32 ± 0.43 for uniform connectivity; mean ± SD; p=−4.8 · 10^−5^, t(158)=−4.18, two-sided t-test).

When we compared the homogeneous and the heterogeneous network, we actually did two comparisions at once: One between homogeneous and heterogeneous connectivities (same or different numbers of connections between different cell types) and another one between a high and low overall number of connections (10% in the homogeneous network vs. about 20% in the heterogeneous network constrained by data). Thus, we need to check which of the two differences sets apart the two types of networks. To this end, we conducted one set of simulations where connectivity was heterogeneous, but scaled to 10% and another one where we set all connectivities to 20%. In both of these cases, persistent activity could be achieved. Thus, we conclude that an increased number of connections is only harmful for persistent activity if they increase the overall number of inhibitory connections.

### Persistent vs. spontaneous activity

In the previous section, we reported conditions under which a biologically tightly constrained network model [21] can exhibit persistent activity. An important goal of the study that introduced this model [21] was to mimic the asynchronous-irregular (AI) background activity [32, 33], which has been observed both in models [32–34] and experiments [33, 35]. Thus, it needs to be tested whether AI activity can be generated in a network that fulfills the constraints for persistent activity (note that we concentrate on the irregularity in the following). We implemented all three conditions (homogeneous short-term plasticity, interneuron parameter variability at 5% of its original value and uniform connectivity at 10% for all cell types) in the original network and assessed both persistent activity and measured the coefficient of variation (*C_V_*), a measure of inter-spike time variability. As expected, persistent activity robustly emerged in this network (Figure 6A) after decreasing the background current from 500 to 120 pA. However, activity was also much more regular compared to the original, heterogeneous network (mean *C_V_* = 0.46 before and mean *C_V_* = 0.14 after the stimulus in Figure 6A compared to 1.04 ± 0.33 in the original network [21], mean ± SD). While removing the constraints on variability of cellular and short-term plasticity parameters did not change the regularity of the activity, using the original, heterogeneous connectivity did: The mean *C_V_* reached 0.64, which is above the lower bound of the values found in the original network. However, persistent activity was not possible in this configuration: Activity levels quickly decay back to baseline once the stimulus is switched off (Figure 6B). This could not be changed by using a range of different background currents and synaptic weight strengths (see below). There are at least two possible explanations for this observation: Persistent activity could either be destabilized by noise, or the dynamic regimes that bring about persistent activity and which generates noisy, irregular behavior may be mutually exclusive. In the following to sections, we present tests of both hypotheses. We found that persistent activity is robust to noise, but that noise indeed needs to be generated at a different synaptic configuration. Finally, we tested to which degree a “noise network” generating irregular activity could overlap with the “signal network” generating persistent activity, both in terms of pyramidal cells and interneurons.

**Fig 6.**
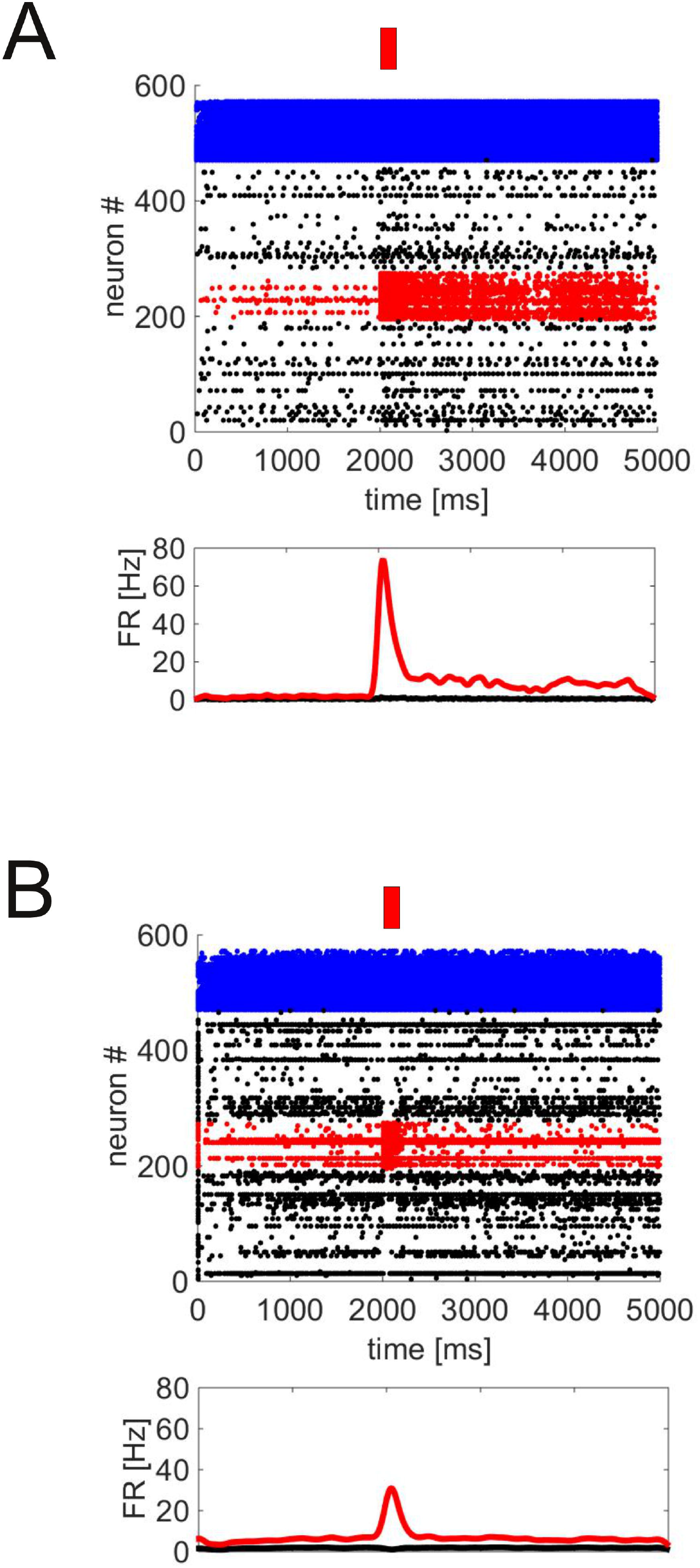
Persistent and spontaneous activity. A: The full network with interneuron cell parameter, connectivity and short-term plasticity heterogeneity removed shows persistent activity, but at low variability. B: When the original, heterogeneous connectivity pattern is used, persistent activity breaks down, but irregular activity emerges. Both panels show raster plots and instantaneous firing rates as in Figure 1A.

#### Robustness against external noise

To investigate the robustness of persistent activity against noise, we added a second set of 1000 neurons to the network and configured the first subnetwork to generate persistent activity (“signal network”, uniform connectivity at 10%) and the second one to generate spontaneous activity (“noise network”, original connectivity from [21], i.e., many fewer connections among pyramidal cells compared to interneurons and compared to pyramidal-interneuron connections). Pyramidal cells from the noise network project onto the pyramidal cells in the signal network with the same synapse parameters as within the noise network, but with uniform connection probabilities *c* between zero and one (which constitutes a minimal perturbation of the signal network). Thus, the two networks simulate spatially separate columns that communicate via pyramidal cells only (see sections below for more interleaved networks). Figure 7A shows the *d*_PA_ values for different values of the coupling constant *c* (only for those simulations which show successful persistent activity, i.e., *d*_PA_ ≥ 0.3). These values do not systematically change as *c* is increased, which shows that persistent activity can survive considerable levels of externally generated noise, compatible with previous results [25]. *C_V_*s within the cell assemblies are consistently above one, largely independent of the values of *c* (1.16 ± 0.06, mean ± SEM over all *c* values). Outside the cell assembly, however, *C_V_*s are strongly modulated by the connection strength to the noise network. Figure 7B shows the *C_V_* both before and after the stimulus presentation as a function of *c* (same simulations as in panel A). *C_V_* increases with c in both cases. However, only *C_V_*s measured after the stimulus presentation, i.e., during persistent activity itself, reach values that are within the range of experimentally measured *C_V_*s *in vivo* (shaded region in Figure 7A, spanning one standard deviation around the mean *C_V_*s measured in a multi-item working memory task in rats [21, 36]). Activity before stimulus presentation is much more regular: *C_V_* values do not reach the in vivo range for any of the *c* values used (stronger connections with *c* > 0.25 ignite persistent activity independent of the stimulus). Apparently the additional (noisy) drive from persistent activity is needed to fuel sufficient activity in the neurons outside the cell assembly, so the noisy activity is a joint product of the noisy dynamics of persistent activity and the synaptic bombardment from outside the network.

**Fig 7.**
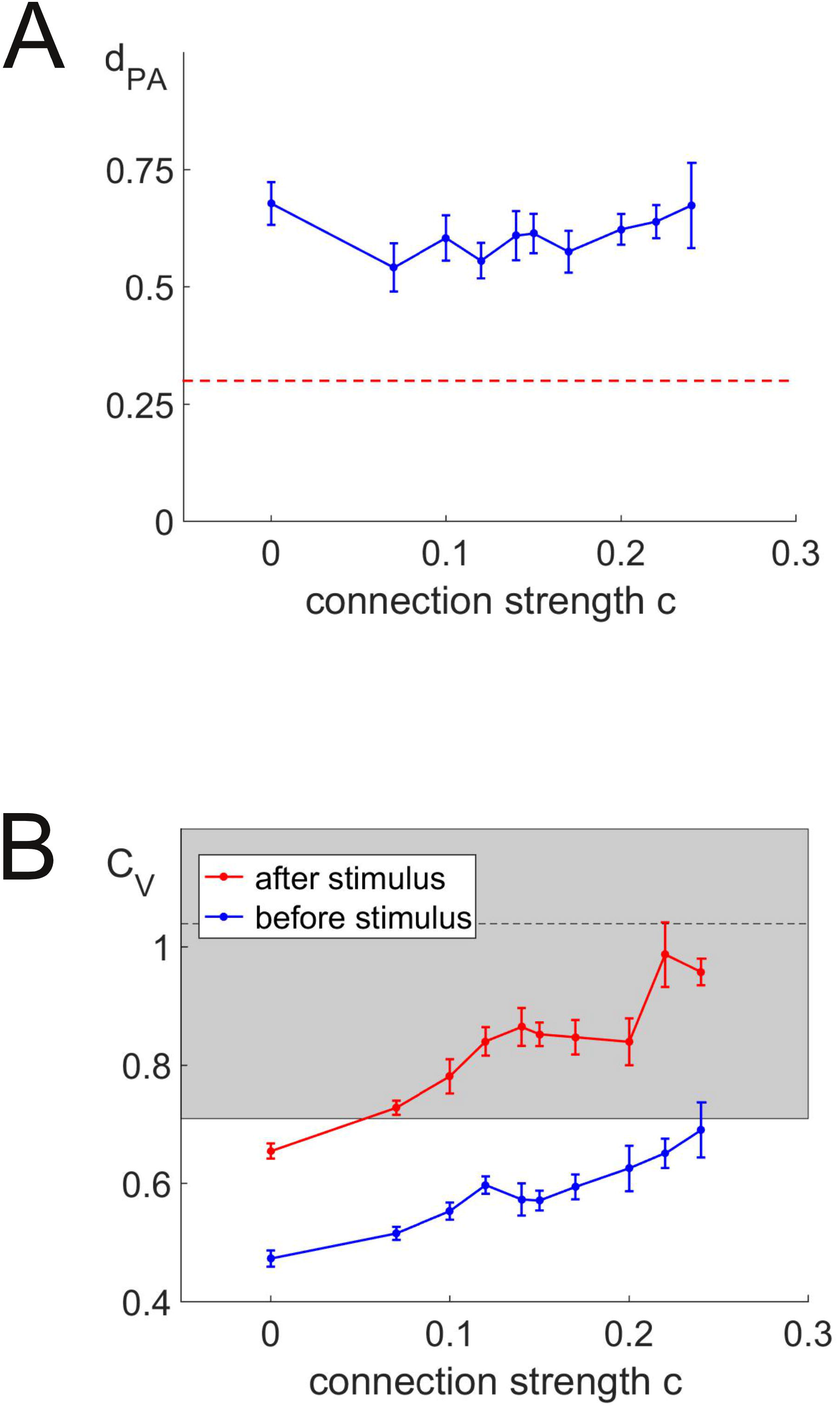
Generation of spontaneous and persistent activity in two coupled networks. A: Persistent activity measure *d*_PA_ as a function of the connectivity *c* (see text for details). B: *C_V_* measured during 500 ms before (spontaneous activity, blue) and after (persistent activity, red) stimulus presentation as a function of the connectivity *c* of the signal network to the noise network (same simulations as in Panel A). The shaded area denotes the range of one standard deviation around the mean (dotted line) of *C_V_* measured experimentally in vivo [21].

In summary, input from the noise network is effective for increasing the *C_V_* of the signal network without harming its ability to generate persistent activity for moderate connectivity between the two networks.

#### Dynamic generation of persistent and spontaneous activity

To answer the question of whether noise and persistent activity can be dynamically generated within the same network (potentially at different times and states), we tried to transform the signal network into a noise network (i.e., a network exhibiting *C_V_*s near one outside the cell assembly) while keeping its ability to generate persistent activity. If this is not possible, the noise within a network exhibiting persistent activity may either have to be generated by external input or by modulation of its properties. As the key difference between the signal and the noise network is in the connection probabilities, we changed these probabilities in the signal network from the uniform 10% back to the original values of the noise network (see previous section) and tried to retrieve persistent activity by compensating for these changes using the synaptic weights: Whenever a connections probability was decreased, we increased the weight of the remaining synapses, and vice versa. Note that this procedure is not intended to mimic a real biological process, but rather to test whether it is possible to construct a network which is capable of both persistent and noise-like activity (see section “Working memory and irregular activity” for a possible way to switch between persistent and irregular activity using neuromodulation). Indeed, we obtained persistent activity with the original connectivity in this way (Figure 8A), but only with fine-tuning of each synaptic weight up to the tenth of a percent. More importantly, this fine-tuning did not yield realistic, noisy activity in the neurons outside the cell assembly: *C_V_*s are very low (mean *C_V_* = 0.27), so the ability to generate noisy activity could not be recovered in a network with persistent activity.

**Fig 8.**
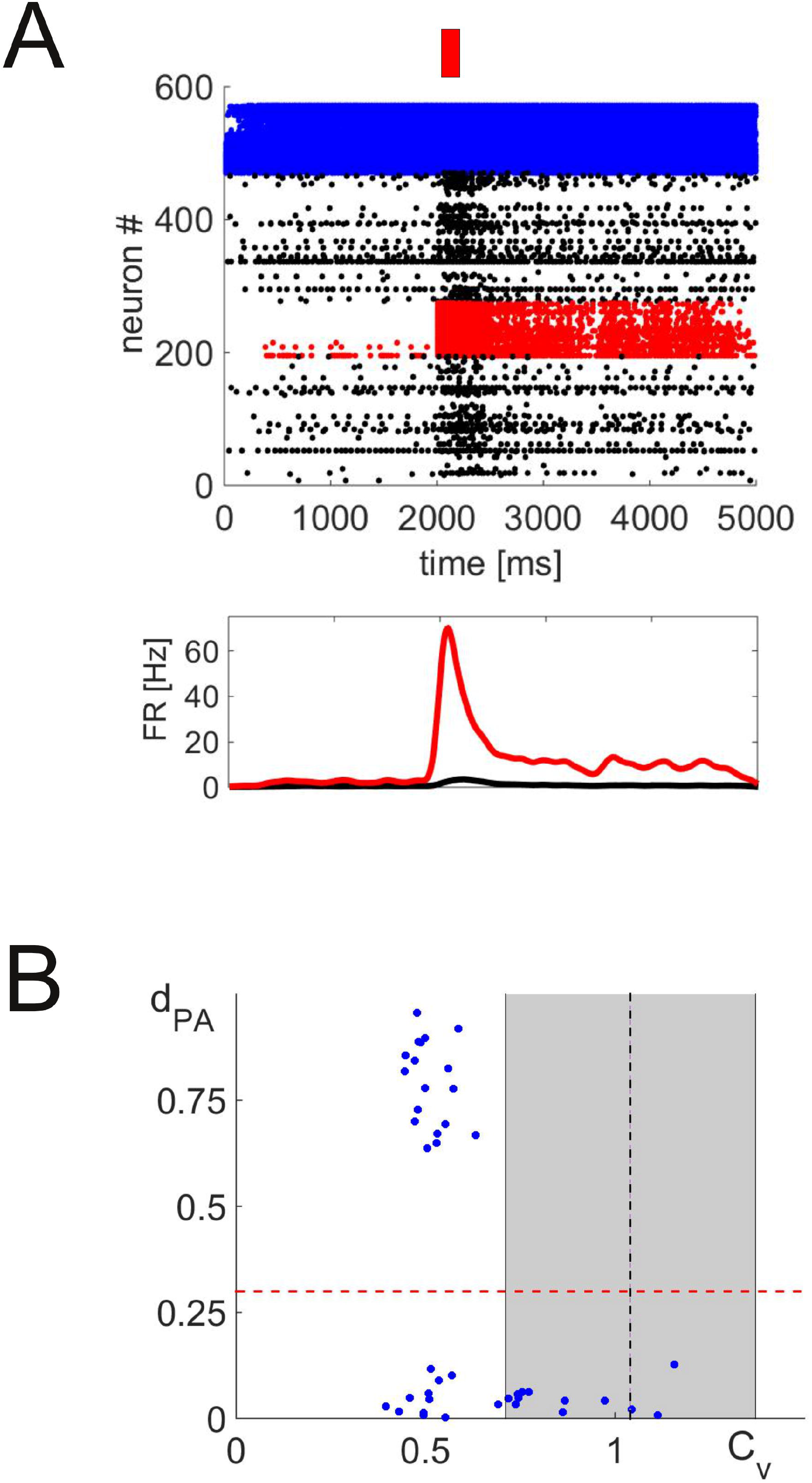
Failure to generate spontaneous and persistent activity in the same network. A: Example of persistent activity with original connectivity values and fine-tuned synaptic weights, but regular activity (*C_V_* = 0.27, raster plot and instantaneous firing rate as in Figure 1A). B: Persistent activity measure *d*_PA_ as a function of the *C_V_* for all attempts to combine spontaneous and persistent activity generation in the same network. The shaded area denotes the range of one standard deviation around the mean (dotted line) of *C_V_* measured experimentally *in vivo*.

We summarize all attempts to combine persistent and spontaneous activity by plotting the mean *C_V_* and *d*_PA_ values in Figure 8B, excluding those configurations which either put the network in an epileptiform state (mean firing rates ≥ 20 Hz for neurons outside the cell assemblies) or yield almost no firing at all (mean firing rates ≤ 0.1 Hz). One clearly sees that successful persistent activity (*d*_PA_ ≥ 0.3, dotted red line) can only be found at low *C_V_* values between 0.5 and 0.6, which are significantly below the Poisson-like value of one usually reported *in vivo*. In particular, all these values are outside the range of *C_V_*s obtained from our reference data set (shaded region in Figure 7B and Figure 8B [21, 36]).

#### Admissible overlap between signal and noise network

As subnetworks in the cortex do not exist in isolation, we next studied to which degree the signal network (generating persistent activity) and the noise network (generating noisy, spontaneous activity) can be mutually interconnected without harming persistent activity, as opposed to the one-directional projection from noise to signal network we described above. We considered two different types of connections (Figure 9A): First, synapses between the two networks, but within the same cell type, i.e. from pyramidal cells of the noise network to pyramidal cells of the signal network, and the same for interneurons (with a connectivity of a fraction *c_P_* of the connectivity within the noise network). And second, synapses between the pyramidal cells of one network and the interneurons of the other (connectivity varied as a fraction *c_C_* of the noise network connectivity). We found that persistent activity can be generated simultaneously with spontaneous activity for arbitrary values of *c_P_*, while even small non-zero values of *c_C_* completely abolished persistent activity (Figure 9B).

**Fig 9.**
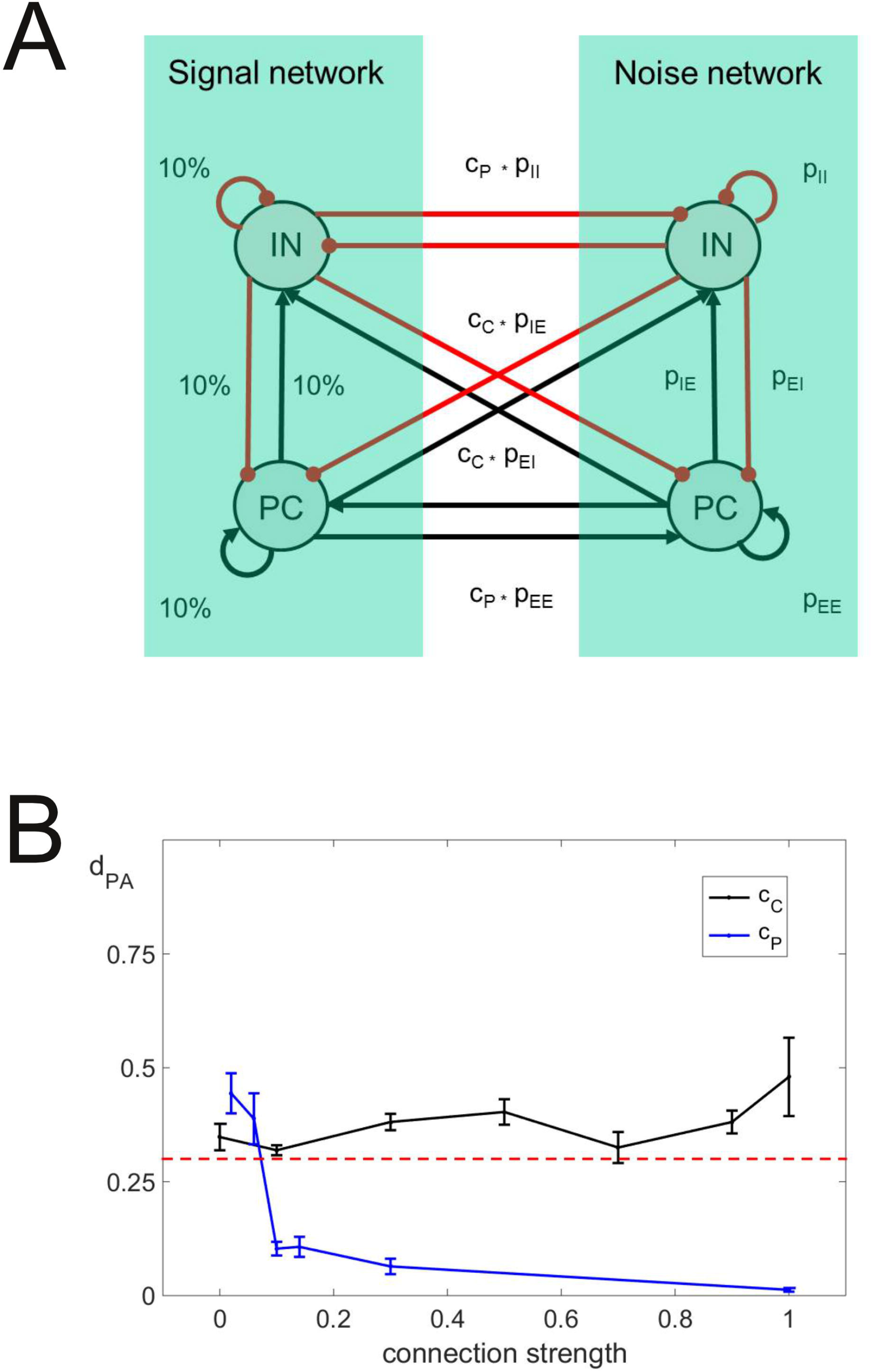
Admissible overlap between signal and noise network. A: Illustration of the network connectivity. A signal network (uniform connectivity of 10%) and a noise network (original connectivity) are connected at variable fractions of the original connectivities. Pyramidal cells and interneurons in both networks are mutually connected at a fraction *c_P_*, and pyramidal cells in one network and interneurons in the other are connected at a fraction *c_C_*. B: Persistent activity measure *d*_PA_ as a function of the connection strengths *c_C_* (black curve) and *c_P_* (blue curve). As one of the strengths is varied, the other one is kept to zero.

From the previous results of this paper, we hypothesized that persistent activity is destroyed by the variability of the inhibitory input from the noise network onto the cell assembly members. To test this hypothesis, we varied the strength of the synaptic input from the interneurons of the noise network to the pyramidal cells of the signal network and compared the statistics of the resulting synaptic conductances (at *c_P_* = 0.3) onto the members of the cell assembly with those conductances emerging without crosstalk between the two networks (*c_P_* =0). We found that the inhibitory currents at *c_P_* = 0.3 and full synaptic strength of the corresponding synapses are higher (average conductance 18.24 ± 1.07 nS for *c_P_* = 0.3 compared to 12.27 ± 0.64 nS for *c_P_* = 0.0; mean ± SEM; p = 2.0 · 10^-45^, t(78)=30.28, one-sided t-test) and more variable (standard deviation 9.23 ± 0.52 nS for *c_P_* = 0.3 compared to 5.90 ± 0.34 nS for *c_P_* = 0.0; mean ± SEM; p = 1.1 · 10^-48^, t(78)=33.90, one-sided t-test) compared to *c_P_* = 0. Only when the weights of the cross-network synapses are decreased as much as 90% of the original value, average strength and variability of the inhibitory inputs are reduced close to the values at *c_P_* = 0 (average conductance 13.76 ± 0.92 nS, standard deviation 6.82 ± 0.50 nS), also restoring persistent activity. Thus, although only 30% of the connections of the noise network project onto the cell assembly members, the corresponding weights must be very strongly decreased to compensate for the effects of the heavy tail in the inhibitory inputs.

### Dynamic mechanism of breaking persistent activity

In this final part of the results section, we investigate the dynamic mechanism of how heterogeneity in the interneuron rheobases breaks persistent activity. This effect seems counterintuitive at first glance, as the stronger input of low-rheobase interneurons should be compensated by the lower input form high-rheobase ones, given a symmetric rheobase distribution. However, inhibitory input to pyramdial cells exhibits two important nonlinearities: If the inhibitory input is too strong, the pyramidal cell ceases firing at all. On the other hand, if the input to interneurons falls below the rheobase, there is no inhibitory input at all, leaving the pyramidal cell constantly active. As heterogeneity of interneuron rheobase increases, more and more pyramidal cells in the cell assembly are being driven into one of these two extremes, taking them out of the bistable regime that is necessary for stimulus-dependent persistent activity.

To show that this mechanism does not depend on the specific configuration of the present model, we use a minimal model of persistent activity [37], comprising only a single pyramidal cell of the simplified Hodgkin-Huxley-type [38] equipped with an GABA and NMDA autapse, which are modelled in the same way as in the present model (conductance-based synapses, NMDA including a magnesium block for low voltages of the postsynaptic cell, see Methods for details). As shown in [37], the positive feedback from the NMDA self-connections enables persistent activity, or more precisely, bistablity between an active state (moderate to high firing rate) and an inactive state (zero firing rate), while the inhibitory GABA self-connection contributes to the stability of the inactive state. We extend this model by filtering the GABA conductances by a number of rectified linear units (ReLU), each mimicking an interneuron with a different rheobase. The slope and the mean rheobase of the ReLU units are chosen to fit the f-I curve of the interneurons in the full model (see Methods for details). Furthermore, we decrease the synaptic condutances to 10% of the values reported in [37] to keep the firing rate persistent activity in a regime similar to the present simulations (up to 80 Hz). This system constitutes a minimal model of a pyramidal cell activating a number interneurons with heterogeneous rheobases, which in turn inhibit that cell, as it is the case of each member of the cell assemblies of the full model considered before.

To analyze the dynamic behavior of the minimal model, we construct a two-dimensional phasespace by computing the instantaneous firing rate FR (inverse of the interspike interval) and the mean input current *I*_syn_ between each two spikes (Figure 10A, following the approach in [39]). In particular, we compute the nullclines of the two variables, curves in phasespace at which the flow in the direction of one of the two variables changes sign, i.e. that variable does not change when starting anywhere the nullcline if the other variable is held constant. The FR nullcline is computed by varying the input current into the cell and compute the average firing rate for each current without synaptic feedback (blue curve in Figure 10A) and the *I*_syn_ nullcline is the mean synaptic current produced at each of these rates (red curve in Figure 10A). The FR nullcline is identical to the f-I curve of the pyramidal cell modelled here and the *I*_syn_ nullcline is the sum of the average NMDA and GABA currents generated at a given firing rate, plus the background current. The geometric shape of the *I*_syn_ nullcline determines the number and stability of fixed points, and thus the ability of the system to exhibit bistability and stimulus-specific persistent activity: Each crossing of the FR and *I*_syn_ nullclines constitutes a fixed point and bistability requires two stable fixed points separated by an unstable one (black dots in Figure 10A). Geometrically, this implies that the slope of the *I*_syn_ nullcline needs to change sign at least once. As the inhibitory input is varied (directly or via rheobase variation), the inhibitory component of the *I*_syn_ nullcline is shifted to the left or to the right. When the inhibitory input becomes too strong, the *I*_syn_ bends more and more to the left, eventually losing the active state fixed point at non-zero firing rates (left green curve in Figure 10B). When the input to the interneurons drops below the rheobase, on the other hand, the inhibitory component of the *I*_syn_ becomes zero and the *I*_syn_ is identical to its excitatory component, which lacks a change in the slope sign and does not cross the FR firing rate at zero firing rate (right green curve in Figure 10B), losing the inactive stater fixed point. In both cases, the system becomes monostable, either in the active or the inactive state.

**Fig 10.**
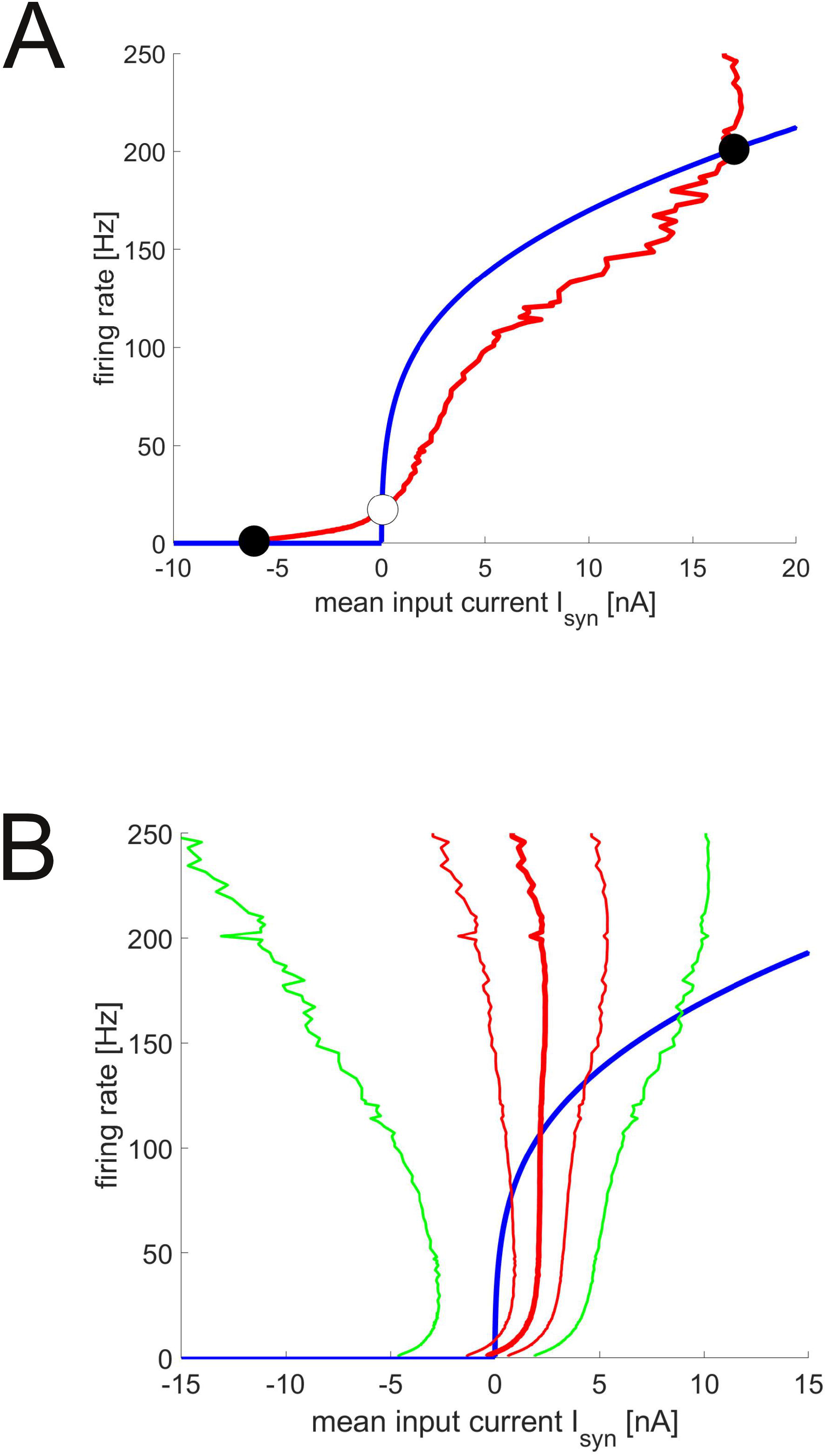
Nullclines of a minimal model of persistent activity. A: Example of nullclines with the three fixed points necessary for bistability. B: Mean (thick red curve), mean plus and minus standard deviation (thin red curves), minimum and maximum (green curves) of the *I*_syn_ nullcline for a large standard deviation of the rheobase of the interneurons (std_rheo=120 pA) together with the firing rate nullcline (blue curve).

To study the effects of heterogeneity in the interneurons, we model a population of 1000 pyramidal cells, each of which receiving a different amount of inhibition and thus being governed by a different *I*_syn_ nullcline (Figure 10B, see Methods for details). As heterogeneity increases, more and more of these cells will be driven out of the bistable range, into one of the two monostable regimes. As a result, the number of bistable neurons, i.e., exhibiting three fixed points, drops with increasing rheobase heterogeneity, which is precisely the effect we observed in the full model using the measure *d*_PA_ (Figure 11A, blue curve). Importantly, changing the (mean) background current does not help to restore bistability, as it simply replaces monostable active by monostable inactive cells or vice versa Figure 11B, compare the red curve with Figure 1C).

**Fig 11.**
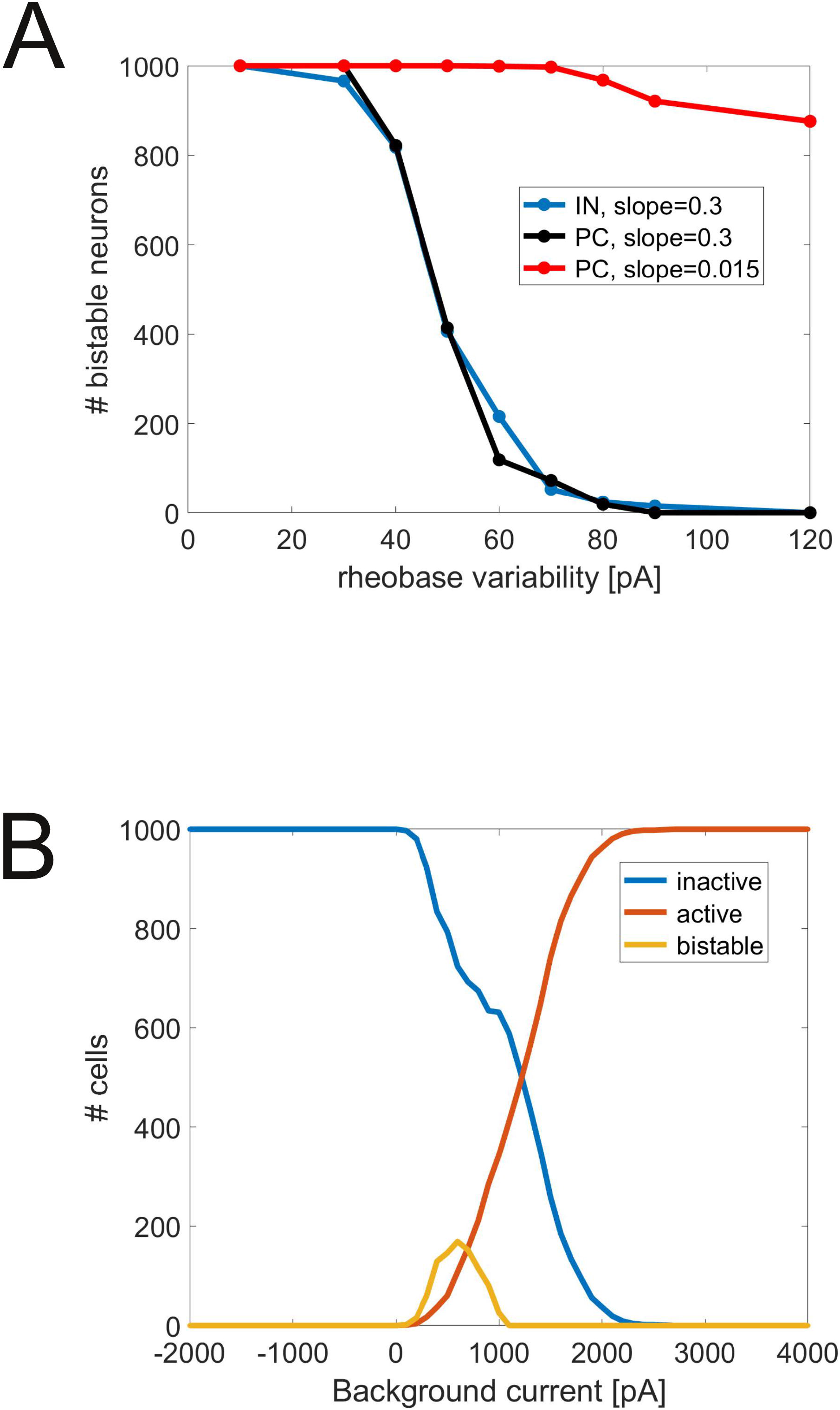
Number of bistable neurons drop as rheobase variability increases. A: Number of bistable neurons, i.e. *I*_syn_ nullclines with three intersections with the firing rate nullcline, as a function of the standard deviation of the rheobase of the ReLU units providing inhibitory input (blue curve) or excitatory input (black and red curves, using two different slopes for the ReLU units). B: Number of bistable (three intersections), active (one intersection at non-zero firing rates) and inactive (one intersection at zero firing rate) neurons as a function of the background current. The standard deviation of the inhibitory ReLU units is set to 60 pA.

The amount of rheobase heterogeneity that is admissible for persistent activtiy depends on the curvature of the *I*_syn_ nullcline, or more precisely, on the range between the current at zero firing rate and the one at the first local maximum where the nullcline changes its slope (e.g. −6 and 15 nA in Figure 10B). This range is largely determined by the strength of the two synaptic inputs. Thus, it is possible to restore persistent activity at a given level of interneuron heterogeneity, e.g., by choosing larger peak conductances. However, such a scaling also increases the firing rate of the active state, illustrated by comparing the (mean) I_syn_ nullclines in Figure 10A and B. Indeed, as we systematically vary the scaling of the peak conductances of the NMDA and GABA currents (Figure 12A), we observe a linear increase in both the maximal admissable rheobase variability (blue curve) and the firing rate (orange curve) as the scaling, and thus, the range of the *I*_syn_ nullcline, is increased.

**Fig 12.**
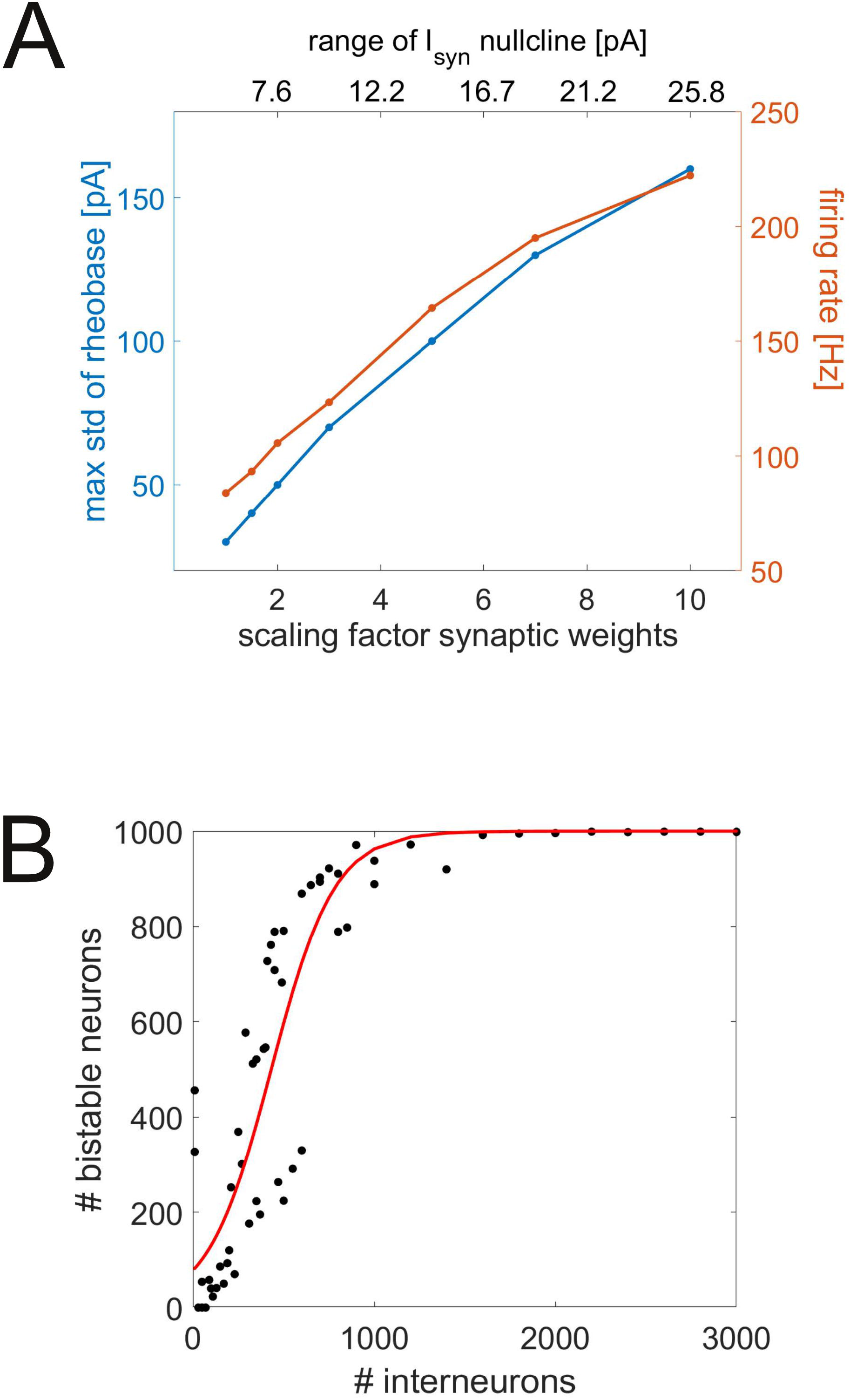
Restoring bistability by increasing peak conductances or number of interneurons. A: Maximum of the rheobase standard deviation without a drop in the number of bistable neurons (blue curve) and firing rate in the active state (orange curve) as both NMDA and GABA peak conductances are being scaled by the same factor. As the weights are scaled up, the range of the (mean) *I*_syn_ nullcline also increases (written above the graph for each value of the scaling factor). B: Number of bistable neurons as the number of interneurons (ReLU units with different rheobase values filtering the inhibitory conductance) is being increased. The standard deviation of the rheobase was set to 60 pA for all simulations.

Next, we compare the effect of interneuron heterogeneity with the effect of pyramidal cell heterogeneity. To that end, we filter the NMDA conductances instead of the GABA conductances by a number of ReLUs, effectively modelling a number of pyramidal cells that are excited by the modelled cell and excite that cell in turn. If the same slope is being used for the pyramdial ReLUs as for the interneuron ReLUs (s=0.3), the number of bistable neurons drop with rheobase heterogeneity in the pyramidal cell in the same way as it did with interneuron heterogeneity (Figure 11A, black curve). However, when mimicking the much shallower slope of the pyramidal cells from the simulations (s=0.05), increasing rheobase heterogeneity to the same extend as in the interneurons does not have a significant effect on the number of bistable neurons, as in the full PFC network (Figure 11A, red curve, cf. (Figure 2A, red curve)). Note that in the full model, pyramidal cell heterogeneity was varied while interneurons exhibited their full heterogeneity, while they are all identical here, explaining the different levels of the curves in the two figures.

Finally, we assess the effect of the number of interneurons projecting on a single pyramidal cell. In all previous simulations, we used 200 ReLU units with different rheobases to compute the inhibitory component for each of the 1000 *I*_syn_ nullclines, each representing a single pyramidal cell. As this number is increased, more and more of the variability in the inhibitory input is averaged out, which leads to a lower standard deviation in the total input to each pyramidal cell. Consequently, the number of bistable neurons increases as the number of simulated interneurons increase (Figure 12B). As apparent in the figure, about 1000 interneurons are sufficient to keep all pyramidal cells in the bistable regime for a rheobase variability of 60 pA.

As the above results suggest that increasing the number of neurons could be a possible way to produce persistent activity in the presence of heterogeneities, we also conducted a number of simulations of the full model with larger networks. As expected, homogeneous networks showed persistent activity, while heterogeneous networks (using the original connectivity) did not using 3000, 5000 or 10000 neurons. Importantly, the 3000 neuron network also showed persistent activity when the rheobase heterogeneity was increased to 10% of the original values, while the 5000 neuron network could even tolerate 50% heterogeneity. However, even the 10000 neuron network was not able to produce peristent activity at full rheobase heterogeneity. So, increasing the network size can in fact be used to extend the range of admissible heterogeneity, but the full range obtained from *in vitro* data can not be averaged out even in a network that is larger than a cortical column (containing about 7500 neurons [40]).

## Discussion

We presented a working memory model based on persistent activity within a biologically validated network model of the prefrontal cortex. We found that the network shows the bistability necessary for stimulus-dependent persistent activity only under the condition that the excitability of the interneurons that inhibit the neurons of the cell assembly is largely homogeneous. The same is true for the strengths of the inhibitory inputs onto the assembly members - a heavy tail in the distribution of these inputs also destroys persistent activity. Furthermore, pairs of neurons need to exhibit the same type of short-term plasticity (if present) for a given combination of cell types. None of these homogeneity criteria is fulfilled in the full network model, nor in the electrophysiological experiments from which the neuron and synapse parameter distributions of the model are taken from [21]. Thus, we argue that persistent activity may require that heterogeneities are compensated by appropriate learning rules such as homeostatic plasticity, at least locally for a given cell assembly.

### Breaking persistent activity by interneuron heterogeneity

We have identified the dynamic mechanism of breaking persistent activity by heterogeneity in the rheobase of the interneurons, namely a spread in the inhibitory input that drives part of pyramidal cells into inactivity and another part into hyperactivity. Using a minimal model of persistent activity, we have shown that this mechanism does not depend on the exact implementation details of the model. Rather, using the language of dynamical systems theory, we have shown that this mechanism is a general dynamic phenomenon that can occur in any system where at least one of the nullclines exhibits enough variability to lead to different number of fixed points for the different components (e.g. pyramidal cells). In this sense, the interneuron rheobase can be seen as a bifurcation parameter that is varied above and below the two bifurcation points between bistability and monostability of the active and inactive state, respectively. In particular, any source of heterogeneity in the inhibitory part of the *I*_syn_ nullcline will have a similar effect, potentially explaining the deleterious effect of heterogeneity in the synaptic weights and in the parameters of short-term synaptic plasticity for persistent activity as well. Also in line with the results in the full network, the minimal model predicts a very limited effect of pyramidal cell heterogeneity (Figure 11A, red curve), which can be explained by the fact that the f-I curve of the pyramidal cells is much shallower compared to those of interneurons (compare red and black curve in Figure 11A).

Using the minimal model, we have identified two potential mechanisms to compensate the effects of heterogeneity in the interneurons: Increasing the synaptic peak conductances and increasing the number of interneurons that innervate a given pyramidal cell (Figure 12). The former is accompanied by an increase in the firing rate (Figure 12A, orange curve). Thus, a low-rate active state is unlikely to be stabilized against interneuron heterogeneity in this way, especially when starting from the even lower weights that are required rates as low as 20 Hz (about 0.5 to 1% of the original weights). In line with these observations, the only simulations of the full model showing peristent activity at full heterogeneity used strongly increased NMDA-to-AMPA ratios and a large cell assembly (e.g. many strong excitatory inputs and lead to epileptiform firing of the entire network at a very high rate.

The latter analysis allows for an estimate of the number of interneurons that are necessary to sufficiently average out their heterogeneity. From Figure 12B, full bistability is recovered starting from about 1000 interneurons (at a rheobase variability of 60 pA, which is close to the value in the full model). Given the number of 7500 neurons in a cortical column [40] and the cell type distributions used in the full model [21], one arrives at 230 local interneurons in L2/3 and and only 40 local interneurons in L5 in a single column, which is clearly not enough to average out rheobase heterogeneities. Taking together all interneuron types, these numbers increase to 780 and 230 in L2/3 and L5, respectively, so the total effect of interneuron heterogeneities may depend on the degree to with these other types of interneurons (e.g. VIP and SOM positive cells) affect persistent activity. A recent study [41] shows that VIP and SOM positive interneurons improves persistent activity rather than hindering it. Finally, we have only considered interneuron heterogeneity in the minimal model, while heterogeneity from other sources will add onto the overall variability of the inhibitory input. In particular, we have seen that the interplay between the (log-normal) distribution of the synaptic weights, the firing rate of interneurons and short-term synaptic plasticity creates a heavy tail of inhibitory inputs (Figure 5). The very strong inputs from that tail often dominate the entire inhibitory input of a neuron, largely irrespective of the number of smaller inputs.

### Working memory and irregular activity

To our knowledge, this study is one of very few to consider the relation between the dynamic generation of persistent activity and irregular activity within the same network (e.g. [34, 42]). Previous working memory models were typically provided with Poisson background spike trains [16, 25, 43], which mimicked the synaptic bombardment with random input from outside the memory network, characterized by coefficients of variation near one and spike-time correlations between input neurons near zero. The generation of noisy activity alone has been subject to many studies (e.g. [32, 44] based on the notion of balanced excitation and inhibition.

The biologically constrained model we use here [21] provides a slightly different mechanism for generating irregular activity, namely a balance between few very active neurons (mostly interneurons) and a vast majority of almost silent neurons (mostly pyramidal cells). While a full analysis of this mechanism is out of the scope of this paper, Figure 5B suggests that irregular activity is being driven by a heavy tail of inhibitory synaptic inputs. Previous studies [42] already suggested that self-sustained irregular activity requires strong synaptic weights. The present study shows that only a few strong inhibitory inputs together with a constant excitatory input also generate irregular activity. In contrast to previous proposals, this form of noise generation is compatible with the frequent observation of a large fraction of neurons that are mostly silent when using recording techniques that are not biased towards spiking such as calcium imaging or *in vivo* patch-clamp (“dark matter theory” of neuroscience, [45–47]).

Importantly, the heavy tail of the inhibitory inputs [22] does not result from the log-normal distribution of the synaptic weights alone, but in concert with heterogeneous firing rates, i.e. strong weights and high firing rates need to coincide to from the tail of the distribution. On the other hand, we have shown that persistent activity breaks down in the present of heterogeneous inhibitory inputs. Thus, at least within our modeling framework, the dynamic basis of irregular activity seems to be incompatible with the more regular, homogeneous synaptic configuration that is needed for persistent activity: A cell assembly needs relatively homogeneous distributions of synaptic inputs to exhibit persistent activity, while heterogeneities in the synaptic inputs are needed to generate irregular activity. The heavy tail explains why the increased inhibitory connectivity in the original network cannot be simply compensated with a decrease in the mean synaptic weights. Such a decrease would only shift the entire input distribution to lower values, but does not abolish the heavy tail that is responsible for most of the increase in the inhibitory conductance.

As a consequence, noisy background activity must be implanted into a cell assembly by its surroundings to show the routinely observed asynchronous-irregular activity [33, 35], while it remains to be investigated whether asynchronicity is also being affected by heterogeneities. Alternatively, the network may be capable of exhibiting both kinds of activity and switch between the signal and noise state by neuromodulation. Dopamine, for instance, is known to stabilize working memory due to a number of modulatory effects on cellular and synaptic properties [25, 48, 49]. Thus, activation of dopamine receptors may compensate the effects of existing heterogeneous inhibitory input. This could enable persistent activity in a noise network, temporarily turning it into a signal network. We can use the minimal model to illustrate how such a modulation could work. In the study [37] from which the model is taken, dopaminergic modulation is modelled by scaling both NMDA and GABA condutances by the same factor, i.e., higher dopamine levels lead to stronger feedback via NMDA and GABA synapses. Thus, we can study the possible effect of dopamine in Figure 12A: As dopamine levels increase, so does the range of the *I*_syn_ nullclines, the maximal admissible standard deviation, and the firing rate. This raises the possibility that high dopamine levels facilitate persistent activity by increasing its firing rates making it more robust against higher levels of interneuron heterogeneity.

When noise is provided from outside the network, as in previous models [16, 25, 43], persistent activity is not impaired (Figure 2B), but shows the irregularity that is seen *in vivo*. Interestingly, *C_V_*s are higher during persistent activity compared to spontaneous activity (Figure 7A), likely due to the additional (noisy) drive provided by the cell assembly neurons to those outside the assembly, which is consistent with observations *in vivo* [50].

Regarding interactions of signal and noise networks, we have shown that inhibitory input from a noise network disrupts persistent activity in a signal network, while excitatory input does not. Interestingly, a recent *in vivo* study has shown that optogenetic activation of somatostatin (SOM)- and parvalbumin (PV)-positive interneurons disrupt persistent activity during the delay period of a memory-guided behavioral task, while vasoactive intestinal peptide (VIP)-positive interneuron activation enhanced task performance [41]. PV interneurons are commonly associated with fast-spiking interneurons, SOM interneurons with Martinotti cells and VIP cells with bitufted and other cross-layer projecting interneurons. The first two types may project beyond a single cortical column, while the latter is largely confined within its own column. The current study may explain the differential effect of VIP and PV/SOM cells on persistent activity under the assumption that signal and noise networks are spatially separated in different cortical columns: Activating PV and SOM cells may elicit inhibitory effects that travels across networks, contaminating the signal networks supporting the delay phase activity. VIP cells, on the other hand, mostly inhibit cells within one column, so there is much less inhibitory crosstalk between different networks. On the other hand, a recent study [51] suggested that cell assembly formation may not be limited to pyramidal cells, but local interneurons can be recruited into the assembly, while far-reaching interneurons such as SOM cells decorrelate from the assembly. Thus, it seems possible that a cell assembly may learn to separate itself from direct inhibitory input from the outside that is harmful to persistent activity, even in the absence of spatial separation.

### From *in vitro* constraints to *in vivo* activity

This study demonstrates a number of constraints on persistent activity within our model, namely homogeneity of interneuron excitability, connection probability and synaptic dynamics within a cell assembly, which are unlikely to be fulfilled if the neurons within the assembly are randomly selected. Thus, we must discuss whether these constraints are relevant for persistent activity in the real brain, and if they are, how they can be fulfilled by biologically plausible mechanisms.

Regarding the biological relevance, we emphasize that all heterogeneities in this model are directly derived from electrophysiological experiments [21]. Nevertheless, we cannot exclude that the variability of cellular and synaptic parameters may be artificially inflated, e.g., by combining data from different experimental conditions. So in theory, if this artificial variability is larger than the variability due to biological diversity, the breakdown of persistent activity could be an artifact of the inflated variability. On the other hand, our results show that even moderate heterogeneity in the interneuron excitability breaks persistent activity (20-25% of the original variability, c.f. Figure 2). Thus, potential artificial variability would have to be inflated at least four-fold over biological variability to break persistent activity on its own, which is highly unlikely. Furthermore, we have drawn all neuron parameters from a multivariate distribution which respects the correlations between these parameters (see Methods for details), so we can exclude that the effect of heterogeneity in one parameter e.g. on firing rate output is compensated by the heterogeneity in another one by mean of correlations between the two [52]. However, we cannot exclude that such compensatory mechanisms exist between neuron and synapse parameters, as we have obtained the latter from the experimental literature. In fact, such correlations could arise the result of homeostatic mechanisms, e.g., compensating the effect of strongly firing interneurons with lower synaptic weights [52] (see below).

For a biologically plausible mechanism to fulfill the homogeneity constraints, we consider that cell assemblies are assumed to be formed by long-term neural plasticity as a result of repeated common input to a set of neurons. This Hebbian plasticity may be complemented by homeostatic plasticity rules that are known to scale synaptic efficacies such that a given average firing rate in the postsynaptic network is maintained [23, 24]. Here, we have considered a (simplified) scaling of the excitatory input into interneurons, such that the differences in excitability are compensated by opposing differences in the input. Another way to limit the effect of the skewed excitability distribution would be to leave the firing rates of the interneurons unaffected, but scale the inhibitory synapses onto pyramidal cells. Recent studies have suggested that both types of homeostatic plasticity may work together to maintain a balance of excitation and inhibition [24], at a spatial resolution that allows fine-tuning of individual synapses [53]. While such a homeostatic mechanism is unlikely to eliminate all heterogeneity in effective excitability, our results suggest that a decrease to a maximum of approximately 20% of the original heterogeneity will be sufficient to enable persistent activity.

Using homeostatic plasticity on the inhibitory synapses could also help to attenuate heterogeneities in the synaptic input: Pyramidal cells that are subject to very high total inputs from interneurons will fire much less than others, leading to a downregulation of its inhibitory weights. Similarly, combined homeostatic plasticity on excitatory and inhibitory synapses could assimilate inputs from synapses with different types of short-term plasticity: On average, facilitating synapses have a stronger total input on the postsynaptic neuron compared to depressing or mixed-type synapses. As an effect, neurons will exhibit higher or lower firing rates, depending on which type of plasticity dominates, and thus, the respective weights will be adjusted to compensate existing heterogeneities.

### Functional benefits from cellular heterogeneity

At first glance, the constraints outlined above appear to limit the utility of cell assemblies as the basis of working memory. However, they could also provide a specific benefit: If each cell assembly is controlled by inhibition of a well-defined strength and timing, the heterogeneity in these parameters between cell assemblies could make each assembly most reactive to a stimulus with specific tuning properties. More specifically, interneuron excitability is an important determinant of the specific frequency of PING-type rhythms (pyramidal-interneuron network gamma [54]) that are generated within a given subnetwork. A cell assembly with a given, mostly homogenous set of interneurons will likely exhibit a rhythm at a narrowly defined frequency and also resonate to input of that particular frequency. If different assemblies exhibit different resonance frequencies, this could help multiplexing multiple memory items at a time, avoiding interference between the assemblies. This proposal is supported by a recent study in monkeys [55], reporting brief gamma bursts during the delay period. Each of these bursts has a well-defined frequency, which varies considerably among different bursts. As cortical oscillations are strongly shaped by interneuron properties, these results are compatible with the existence of a spectrum of cell assemblies, each of which controlled by a relatively homogeneous interneuron population.

### Model predictions

The current model makes a number of concrete predictions which could be tested experimentally. Here, we make them explicit and briefly discuss methodological issues of this test.

1. Interneurons within a single cell assembly exhibit similar firing rates.
2. Isolated networks that show persistent activity have more regular spontaneous activity (lower coefficient of variation) before stimulus onset compared to those in which activity does not persist.
3. Synaptic connections between the same cell types within a cell assembly have the same type of short-term synaptic plasticity (namely, facilitating for connections among pyramidal, depressing for interneuron-to-pyramidal connections or those among interneurons).

All three predictions require an experimental identification of cell assemblies and a subsequent analysis of cellular or synaptic properties. A number of computational methods are now available for cell assembly detection in electrophysiological data, the most recent one even spanning a range of lag times [56]. The non-trivial process of detecting these assemblies may be one of the reasons why the above properties have not been identified before: Assemblies are not necessarily spatially localized, thus anatomical and physiological recordings that are focused in a small spatial volume may fail to see the organization of cellular and synaptic properties within a cell assembly. If these recordings are combined with a detection of the cell assembly, though, one could test, e.g., the first prediction by comparing the firing rates of interneuron units. This could be done both *in vivo* and *in vitro*. Prediction 2 requires the comparison of activity in the same units before and after a stimulus and the isolation of the memory network from most of its environment, both of which should be possible *in vitro*. Finally, prediction three concerns the dynamic properties of synapses within an assembly. Organotypic slices provide access to these properties, with a good chance of intact assembly structures, although it is not always clear to what extent *in vitro* results can be transferred to in vivo networks.

Naturally, the predictions rest on the assumption that the network model [21] provides a reasonably valid image of the prefrontal cortex, despite the necessary simplifications, such as a specific selection of cell types and cortex layers, a phenomenological model of the neurons [28] and a constant background current for all pyramidal cells and interneurons of a given layer. We have previously shown that the neuron model provides an accurate description of neuron’s responses to fluctuating currents [28] and the network model veridically reproduces many key features of baseline *in vivo* activity [21]. Nevertheless, the model needs to be further tested in terms of its ability to reproduce a wider range of dynamic regimes and functions of the prefrontal cortex.

### Heterogeneity in previous working memory models

While there is a wide range of models of working memory based on persistent activity, most of them use homogeneous parameters and few have investigated the effects of heterogeneities on working memory performance; to our knowledge, there are four studies on variants of the ring model [17–20] and another one on a related parametric memory model [57], investigating the effect of random and spatially structures heterogeneities on working memory. In the ring model [16], neurons are spatially organized on a ring and form the strongest connections to their nearest neighbors, effectively forming a continuum of cell assemblies along the ring. Thus, input at any point of the ring can cause a persistent bump of activity in the neurons nearby. Random heterogeneities, e.g., in neural excitability, cause these bumps to drift towards specific points on the ring, compromising their ability to encode the spatial position of the input over time [17–20]. The reason for this drift is the fact that the energy landscape along the ring is basically flat (neutral stability on the ring), such that any point of the line can be represented. Random heterogeneities create hills and valleys in this landscape, so activity drifts towards its local minimum. Homeostatic plasticity has shown to effectively flatten out these local minima such that the representation becomes stable again. In contrast to random heterogeneities, spatially structured heterogeneities are deliberately put into the networks to fulfill a specific function [20, 57], in one case even to counteract the harmful effects of the random heterogeneities [20].

In the present model, we focus on random heterogeneities in the cell parameters, which destroy working memory when present in the interneurons. The cause of this problem is very different from the one in the ring model: Instead of shifting persistent activity into some uninformative region, it is either completely lost or permanently activated without a stimulus (c.f. Figure 1) - in short, the system loses its bistability. In the ring model, the instability was compensated using either homeostatic plasticity targeting the activity of pyramidal cells [17], short-term plasticity [18, 19] or a systematic structure of the energy landscape, effectively segmenting the line attractor into a discrete set of attractors [20]. In principle, however, this instability could be removed by considering a ring with a sufficiently large number of neurons, which would statistically equalize populations representing different spatial positions. The loss of bistability due to the skewed distribution of interneuron excitability, on the other hand, must be compensated for persistent activity to work at all. These considerations emphasize the need to work with parameter distributions that are derived from experimental studies.

### The current state of persistence-based working memory models

Persistent activity in cell assemblies in the prefrontal cortex has long been the dominant model for the mechanism of working memory. Recently, this dominance is challenged by a a number of alternative model frameworks which are based, e.g., on short-term synaptic modifications [15], dynamic activity patterns [11], or short-lived attractor states [58], potentially paced by beta, gamma and theta oscillations [59]. The theoretical proposals are accompanied by experimental observations that delay activity may be more complex than simply maintaining neurons at elevated firing rates [11, 55]. Supporters of persistence-based models counter these arguments showing that the alternative models “could mediate only a limited range of memory-dependent behaviors” and are not mutually exclusive with stimulus-dependent persistent activity [6].

We argue that tests of the current model’s predictions would also contribute to the evidence in favor or against the general concept of persistence-based working memory models. As the role of oscillations in working memory has been addressed in a number of recent studies [55, 60, 61], we note that we also see slow, quasi-periodic interruptions of persistent activity in our model (e.g. Figure 1B, frequencies in the delta range, 0.6-2 Hz), and faster (beta/gamma) oscillations can by induced by strongly decreasing the background current into the interneurons while increasing the pyramidal-to-interneuron connections, leaving interneuron activity to be more controlled by pyramidal cell activity and thus enabling a PING-type rhythm (pyramidal-interneuron network gamma [54]). As outlined above, this would lead to a range of assemblies, each of them associated with its own, relatively well defined frequency signature, as seen in [55]. As proposed in [59], increased working memory load would then be associated to more gamma-synchronized assemblies being active at the same time. This would not only result in increased gamma power (e.g. [55, 62]), but also predict that the overall gamma band would be more smeared out, because it would result from several assemblies oscillating at different frequencies. All these proposals can now be tested in a biologically validated modeling framework, contributing to our understanding of the neural underpinnings of working memory.

## Methods

### Model description

The data-driven model of the prefrontal cortex used here is described in detail in [21]. Briefly, the model consists of 1000 neurons modeled by a simplified version (simpAdEx [28]) of the adaptive exponential integrate-and-fire neuron (AdEx [63]) which was optimized for high-throughput fitting to in vitro electrophysiological data:

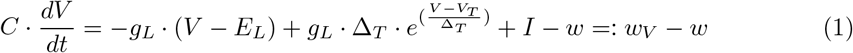

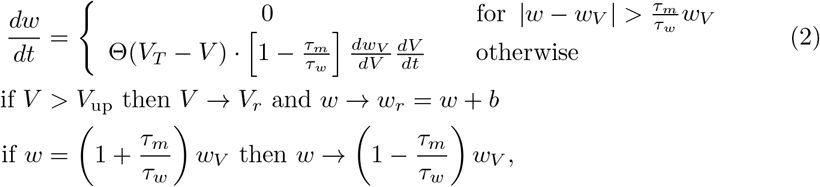

where *C* is the membrane capacitance, *g_L_* a leak conductance (with reversal potential *E_L_*), *τ_m_* and *τ_w_* are the membrane and adaptation time constants, respectively, Θ denotes the heavy-side function, and *w_V_* is the V-nullcline of the system as defined in Equation 1. Like the full AdEx [64], this model consists of one differential equation for the membrane potential *V* (including an exponential term with slope parameter Δ_*T*_, which causes a strong upswing of the membrane potential once it exceeds *V_T_*), and one for an adaptation variable *w*, and can reproduce a whole variety of different spiking patterns [28]. A spike is recorded whenever V crosses *V*_up_, at which point the voltage is reset to *V_r_* and spike-triggered adaptation is simulated by increasing *w* by *a* fixed amount *b*.

Based on about 200 neurons from rodent PFC, we generated multivariate parameter distributions for five different electrophysiological neuron types [21], namely pyramidal cells in layer 2/3 and 5, fast-spiking, bitufted and Martinotti interneurons. These distributions respect the full covariance structure between the eight parameters of model, which was estimated from the joint data set of all fits of the model to each individual neuron. The simulated neurons are embedded in a laminar and columnar network structure that distinguishes pyramidal cells from superficial (L2/3) and deep layers (L5) as well as four subsets of interneurons which either project within the same layer and column (fast-spiking interneurons), or across layers (bitufted interneurons), columns (large basket cells, with the same electrophysiological properties as the pyramidal cells of the respective layer [65]) or both across layers and columns (Martinotti cells). Neurons from each of these subclasses are randomly connected with different connection probabilities. These values, as well as the statistics on synaptic weights (following a log-normal distribution [66]) and connection delays are directly obtained from the experimental literature. Neurons are connected through conductance-based AMPA-, GABA_A_-, and NMDA-type synapses, with kinetics modeled by double exponential functions [39], which time constants again taken from the literature (see Figure SS1 Fig for a schematic of the network architecture and Tables SS1 Table to SS4 Table for a list of the network parameters).

Synapses are equipped with short-term plasticity dynamics implemented by the corrected version [67] of the Tsodyks and Markram model [29]

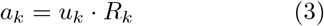

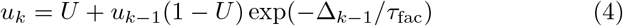

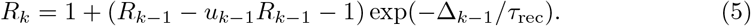

These recursive equations describe the dynamics of the relative efficiency *a*(*t*_sp_k__) across series of spikes, with initial conditions *u*_1_ = *U* and *R*_1_ = 1, where *t*_sp_k__ is the interval between the (*k* – 1)th and the *k*th spike. Model parameters *U*, *τ*_rec_ and *τ*_fac_ were specified according to [30] and [31] who differentiated between facilitating, depressing or combined short-term dynamics, for both excitatory and inhibitory connections. The cell types of the pre- and postsynaptic neurons determine which of these classes is used for each individual combination. In particular, connections among pyramidal cells, among interneurons and between fast-spiking interneurons and pyramidal exhibit all three types of connections to varying degrees (pyramidal-pyramidal: 45% E1, 38% E2, 17% E3 [31]; interneuron-interneuron: 29% E1, 58% E2, 13% E3 [30], fast-spiking interneuron-pyramidal: 25% E1, 50% E2, 25% E3 [30]). All neurons are driven by constant background currents. The values of these currents are the only parameters which are not directly obtained from the in *vitro* literature, but are estimated in a self-consistent way from the network activity itself (see [21] for details).

Importantly, the model has been validated with neuronal data on two levels. On the single-cell level, the fits obtained from the simpAdEx model were tested with in *vitro* data which are fundamentally different from those used for the fit [28], namely fluctuating currents generated by filtering a Poisson spike train with the synapse model described above, as opposed to standard constant DC currents which were used for fitting. The responses of the model neurons to the fluctuating currents matched those of the real neurons within the bounds of their own reliability [28]. On the network level, we compared the activity of the model network which the stationary extracellular recordings from rats *in vivo* during a multi-item working memory task, and intracellular recordings from anesthetized rats. We assessed spike train statistics such as mean, *C_V_*, autocorrelation of interspike intervals and cross-correlations between interspike intervals from pairs of neurons, power spectra of local field potentials and the distribution of membrane potential fluctuations. In almost all of these measures, the distributions from the experimental recordings and the model network were statistically indistinguishable.

### Implementation of cell assemblies

For a given simulation, a random population of 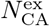 pyramidal cells and 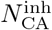 interneurons was defined as a cell assembly. Synaptic connections within an assembly were strengthened by either increasing all peak conductances (synaptic weights) of existing connections by a factor *s*_CA_ or by rewiring the neurons with connection probabilities increased by a factor *p*_CA_. Unless otherwise stated, 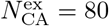 and 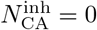 was used. For graphical representations in raster plots, all members of a cell assembly are grouped together in the middle of the respective neuron pool.

### Stimulus presentation

External stimuli were simulated by Poisson spike train with firing rate *f*_inp_ for period of *T*_inp_ ms in a number of *N*_inp_ excitatory input neurons. Two cases are considered: In the first case, a single input neuron (*N*_inp_=1) is randomly connected to L2/3 cells, using the connection probability for pyramidal cells within that layer. In the second case, there is one input neuron for each pyramidal cell in the assembly 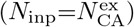, with connections from each input neuron to one of the assembly neurons. In both cases, all synaptic peak conductances of the input are set to *s*_inp_ times the peak conductance of excitatory synapses within the network. Unless otherwise stated, the following values are used for these parameters: *f*_inp_ = 1000 Hz, *T*_inp_ = 50 ms and *s*_inp_ = 1.

### Measures of persistent activity

To quantify to which degree a cell assembly exhibits persistent activity, we define the measure *d*_PA_ as follows

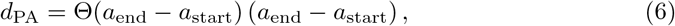

*a*_end_ and *a*_start_ are normalized measures of the activity during 500 ms before the stimulus and the last 500 ms of the simulation, respectively. Θ is the heavy-side function, setting *d*_PA_ to zero if activity is higher in the beginning (i.e. *a*_end_ – *a*_start_ < 0). The activity measures are computed by comparing the firing rate in the cells of the assembly with those outside the assembly. An assembly cell is regarded as activated if its average firing rate during the observed time interval exceeds those of the average firing rate of all non-assembly (pyramidal, layer 2/3) cells. Normalized activity *a* is then defined as the fraction of activated cells in the assembly. Thus, by definition, *d*_PA_ ranges between zero and one. The maximum of one is reached when all cells in assembly are activated at the end of the simulation and none of them before the stimulus, which reflects the desired case of stimulus-induced persistent activity. On the other hand, *d*_PA_ is zero if either none of the assembly cells is activated in the end (decayed activity) or more cells are activated before the stimulus (*a*_end_ – *a*_start_ < 0, spontaneous persistent activity).

### Variation of model parameters

We systematically varied a number of parameters of the model in order to find configurations allowing for persistent activity. In particular, we varied the size of the cell assembly between 30, 50, 80 and 470 neurons (the latter being the number of neurons in L2/3), the ratio between NMDA and AMPA conductances between 1, 2, 3, 4, 6 and 8 times the original value, the peak conductance among pyramdial cells between 1, 5, 10, 15 and 20 times the original value, the scaling factor *s*_CA_ of the synaptic weights within the cell assembly between 1, 5, 15 and 20, connectivity within the cell assembly between 10 and 40% and both background currents *I*_ex_ into pyramidal cells and *I*_inh_ into interneurons between 0 and 1000 pA. We conducted all of these changes in isolation (using a default of 80 cell assembly neurons, the original ratio of NMDA and AMPA conductances as well as the original value of peak conductances among pyramidal cells, a scaling factor for the weights within the cell assembly of 5 and a connectivity of 10% as well as *I*_ex_ and *I*_inh_ values of 300 and 200 pA, respectively) and also assessed a considerable number of these modifications together. Overall, we performed 720 simulations with the full model, summarized in Figure 1C (except for those where the full L2/3 was used as a single assembly, as they exhibit epiletiform activity following stimulus presentation).

### Minimal model of persistent activity

To assess the dynamic mechanisms of persistent activity and the effect of rheobase heterogeneitities in more detail, we employ a minimal model of persistent activity [37] comprising a single pyramidal cell modelled by a simplified version of the Hodgkin-Huxley equations [38] that excites itself by an NMDA autapse and also inhibits itself by a GABA autapse. We modify this model in three ways: First, we decrease the strenghts of the synaptic weights to 10% of the original values. This allows for firing rates (80 Hz) that are closer to the low rates (about 20 Hz) seen in our full simulations (compared to over 200 Hz at the original weights, c.f. Figure 12A). Further decrease of the weights would have required too much fine-tuning to show the principle results, but we also discuss the effect of the scaling in the main text. Second, we filter the inhibitory conductances by a function *G_I_* that mimicks interneuron heterogeneity. This function is described in details below. In some simulations (indicated at the text), we apply a function *G_E_* of the same form to the excitatory conductances, but with different parameters. Finally, we simulate a population of 1000 pyramidal cells instead of one, which only differ in their particular realization of *G_I_* or *G_E_*. This population of models (c.f. [52]) allows to assess the impact of heterogeneity on the dynamics of the pyramidal cell.

The equations governing the dynamics of one of the pyramidal cells are given by:

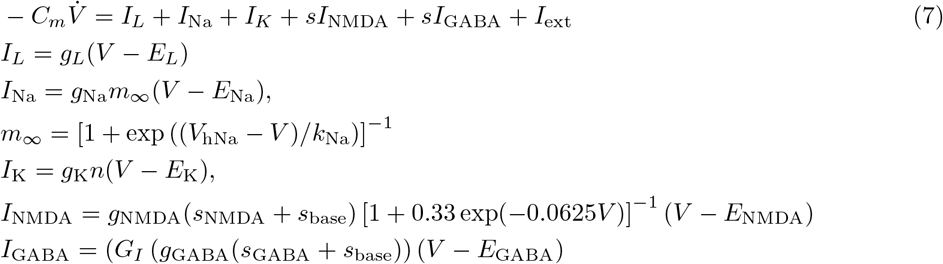

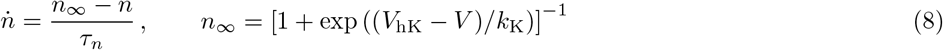

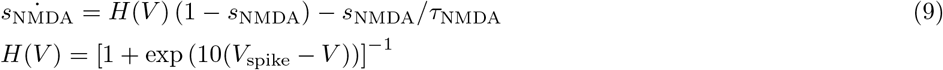

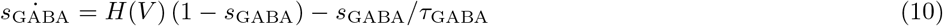

By applying the function *G_I_*, we interpret the inhibitory conductance as an input to a population of interneurons, which are modelled as rectified linear units (ReLU) representing the f-I curve of the interneurons, each with a different rheobase drawn from an normal distribution. The conductance is first converted into an input current, second into a firing rate and third back into a conductance. In this way, one inhibitory conductance is being generated for each interneuron and all these conductances are averaged over all interneurons projecting to the pyramidal cell, which is the output of *G_I_*. The first two steps - conversion of the conductance into a current and into a firing rate - are being conducted in a self-consistent way by finding the intersection between the f-I curve of the current interneuron and a curve describing the current that is being generated at a given firing rate, given the input conductance. That second curve is computed from the numerically obtained relation between firing rate and average membrane potential during the interspike interval of the simulated neuron and the relation between current *I* and conductance g, *I* = *g*(*V* – *E*). Once the firing rate is obtained, it is multiplied with the GABA peak conductance gcABA, which is equivalent to the computation of the average over a long random spike train at that rate, convoluted with the response function of the synapse, given a normalization of the area under that function to one. We tested different variants of implementing *G_I_* and found that results did not qualitatively depend on these details.

The different pyramidal cells in the population differ only in the inhibitory input they receive via *G_I_*. In particular, we randomly pick 20% of the simulated interneurons (200 by default) and construct the synaptic input in *G_I_* only from these 20%.

The phase space of the model is being constructed by considering one pair of subsequent spikes at a time and computing both the instantaneous firing rate FR (inverse of the interspike interval) and the average of the input currents (*I*_syn_ = *I*_NMDA_ + *I*_GABA_ + *I*_ext_) over that interval. The time series of these two variables over all pairs of spikes constitutes the trajectory of the system in the phasespace. The nullclines were being computed by varying the input current into the cell and compute the average firing rate for each current without synaptic feedback (FR nullcline) and the mean synaptic current produced at each of these rates (*I*_syn_). Intersections of the nullclines constitute fixed points of the system and we verified that the trajectories converged to one of these fixed points for different parameter sets.

To classify neurons as bistable, active or inactive, we compute the intersections between FR and *I*_syn_ nullclines for each of the simulated pyramidal cells. Cells with three intersections are labeled as bistable, cells with only one intersection at zero firing rate as labeled as inactive and cells with one intersection at non-zero firing rates are termed active. Importantly, in real, connected network, the number of bistable cells also affects the shape of the *I*_syn_ nullclines: Inactive pyramidal cells in the cell assembly do not contribute to the excitatory feedback and constantly active cells also need to be excluded from the assembly to avoid premature activation before a stimulus arrives. We mimick these constraints by scaling the excitatory peak conductance by the fraction of bistable neurons. After that, the numbers of each types of cells is counted again and the procedure is repeated until convergence to a fixed number of each type (always the case after at most five iterations).

## Acknowledgments

We thank Michelle McCarthy for valuable discussions and continuous support during this research.

## Supporting information

**S1 Fig. Schematic of the network architecture.** (A) Laminar structure of a single network column (only one column is simulated in this study). Arrow widths represent relative strength of connections (black: excitatory, gray: inhibitory), i.e. the product of connection probability and synaptic peak conductance. (B) Left panel: Distribution of three different short-term plasticity types over different combinations of pre- and postsynaptic neuron types. Arrows from or to one of the shaded blocks (rather than from or to a single neuron type) denote connection types that are identical for all excitatory (PC) or inhibitory (IN) neurons. Where all three types are drawn, they are randomly distributed over all synapses between these two neuron types according to the probabilities given in the figure (termed “heterogeneous STP” in the the text). Right panel: Illustration of the postsynaptic potential in response to a series of presynaptic spikes for three types of short-term synaptic plasticity for excitatory (E1 to E3) and inhibitory synapses (I1 to I3). The figure is taken from [21].

**S1 Table. Neuron parameters.** Mean and standard deviation of the parameters of the simpAdEx model for the five different neuron types used in the network (PC: Pyramidal cell, FS: Fast-spiking interneuron, BT: Bitufted interneuron, MC: Martinotti cell). The table is taken from [21].

**S2 Table. Cell numbers.** Relative numbers of cells for each type. PC: pyramidal cell, IN: interneuron, see Materials and Methods section in [21] for interneuron subtypes. The table is taken from [21].

**S3 Table. Synaptic parameters.** Mean ± standard deviation of the parameters of the synapses connecting the different pre- and postsynaptic neuron types (*p*_con_: connection probability, *g*_max_: peak conductance, *τ_D_*: transmission delay). The table is taken from [21].

**S4 Table. Short-term synaptic plasticity.** Mean (standard deviation) of the parameters of the six types of short-term synaptic plasticity. The table is taken from [21].

## References

1. Baddeley A. Working memory: theories, models, and controversies. Annual review of psychology. 2012;63:1–29.

2. Goldman-Rakic PS. The physiological approach: functional architecture of working memory and disordered cognition in schizophrenia. Biological psychiatry. 1999;46(5):650–661.

3. Meyer-Lindenberg A, Weinberger DR. Intermediate phenotypes and genetic mechanisms of psychiatric disorders. Nature Rev Neurosci. 2006;7:818–827.

4. Durstewitz D, Seamans JK, Sejnowski TJ. Neurocomputational models of working memory. Nature neuroscience. 2000;3:1184–1191.

5. Sreenivasan KK, Curtis CE, D’Esposito M. Revisiting the role of persistent neural activity during working memory. Trends in cognitive sciences. 2014;18(2):82–89.

6. Riley MR, Constantinidis C. Role of prefrontal persistent activity in working memory. Frontiers in systems neuroscience. 2015;9.

7. Fuster JM. Unit activity in prefrontal cortex during delayed-response performance: neuronal correlates of transient memory. Journal of Neurophysiology. 1973;.

8. Goldman-Rakic PS. Cellular basis of working memory. Neuron. 1995;14(3):477–485.

9. Curtis CE, Lee D. Beyond working memory: the role of persistent activity in decision making. Trends in cognitive sciences. 2010;14(5):216–222.

10. Passingham D, Sakai K. The prefrontal cortex and working memory: physiology and brain imaging. Current opinion in neurobiology. 2004;14(2):163–168.

11. Stokes MG. ‘Activity-silent’working memory in prefrontal cortex: a dynamic coding framework. Trends in cognitive sciences. 2015;19(7):394–405.

12. Wang XJ. Synaptic reverberation underlying mnemonic persistent activity. Trends in neurosciences. 2001;24(8):455–463.

13. Compte A. Computational and in vitro studies of persistent activity: edging towards cellular and synaptic mechanisms of working memory. Neuroscience. 2006;139(1):135–151.

14. Lansner A. Associative memory models: from the cell-assembly theory to biophysically detailed cortex simulations. Trends Neurosci. 2009;32(2):178–186.

15. Mongillo G, Barak O, Tsodyks M. Synaptic theory of working memory. Science. 2008;319(5869):1543–1546.

16. Compte A, Brunel N, Goldman-Rakic PS, Wang XJ. Synaptic Mechanisms and Network Dynamics Underlying Spatial Working Memory in a Cortical Network Model. Cereb Cortex. 2000;10:910–923.

17. Renart A, Song P, Wang XJ. Robust spatial working memory through homeostatic synaptic scaling in heterogeneous cortical networks. Neuron. 2003;38(3):473–485.

18. Itskov V, Hansel D, Tsodyks M. Short-term facilitation may stabilize parametric working memory trace. Frontiers in computational neuroscience. 2011;5:40.

19. Hansel D, Mato G. Short-term plasticity explains irregular persistent activity in working memory tasks. Journal of Neuroscience. 2013;33(1):133–149.

20. Kilpatrick ZP, Ermentrout B, Doiron B. Optimizing working memory with heterogeneity of recurrent cortical excitation. The Journal of Neuroscience. 2013;33(48):18999–19011.

21. Häss J, Hertag L, Durstewitz D. A Detailed Data-Driven Network Model of Prefrontal Cortex Reproduces Key Features of In Vivo Activity. PLOS Comput Biol. 2016;12(5):e1004930.

22. Buszáki G, Mizuseki K. The log-dynamic brain: how skewed distributions affect network operations. Nature Rev Neurosci. 2014;15(4):264–278.

23. Turrigiano GG, Leslie KR, Desai NS, Rutherford LC, Nelson SB. Activity-dependent scaling of quantal amplitude in neocortical neurons. Nature. 1998;391(6670):892–896.

24. Turrigiano G. Too many cooks? Intrinsic and synaptic homeostatic mechanisms in cortical circuit refinement. Annual review of neuroscience. 2011;34:89–103.

25. Brunel N, Wang XJ. Effects of neuromodulation in a cortical network model of object working memory dominated by recurrent inhibition. Journal of computational neuroscience. 2001;11(1):63–85.

26. Myme CIO, Sugino K, Turrigiano GG, Nelson SB. The NMDA-to-AMPA Ratio at Synapses Onto Layer 2/3 Pyramidal Neurons Is Conserved Across Prefrontal and Visual Cortices. J Neurophysiol. 2003;90(2):771–779. doi:10.1152/jn.00070.2003.

27. Wang H, Stradtman GG, Wang XJ, Gao WJ. A specialized NMDA receptor function in layer 5 recurrent microcircuitry of the adult rat prefrontal cortex. Proc Natl Acad Sci. 2008;105(43):16791–16796. doi:10.1073/pnas.0804318105.

28. Hertäg L, Hass J, Golovko T, Durstewitz D. An Approximation to the Adaptive Exponential Integrate-and-Fire Neuron Model Allows Fast and Predictive Fitting to Physiological Data. Front Comput Neurosci. 2012;6. doi:10.3389/fncom.2012.00062.

29. Markram H, Wang Y, Tsodyks M. Differential signaling via the same axon of neocortical pyramidal neurons. Proc Natl Acad Sci. 1998;95(9):5323–5328.

30. Gupta A, Wang Y, Markram H. Organizing Principles for a Diversity of GABAergic Interneurons and Synapses in the Neocortex. Science. 2000;287(5451):273–278. doi:10.1126/science.287.5451.273.

31. Wang Y, Markram H, Goodman PH, Berger TK, Ma J, Goldman-Rakic PS. Heterogeneity in the pyramidal network of the medial prefrontal cortex. Nat Neurosci. 2006;9(4):534–542. doi:10.1038/nn1670.

32. Brunel N. Dynamics of sparsely connected networks of excitatory and inhibitory spiking neurons. J Comp Neurosci. 2000;8(3):183–208.

33. Destexhe A, Rudolph M, Pare D. The high-conductance state of neocortical neurons in vivo. Nature Rev Neurosci. 2003;4:739–751.

34. Kumar A, Schrader S, Aertsen A. The High-Conductance State of Cortical Networks. Neural Comput. 2008;20:1–43.

35. Renart A, de la Rocha J, Bartho P, Hollender L, Parga N, Reyes A, et al. The Asynchronous State in Cortical Circuits. Science. 2010;327(5965):587–590.

36. Lapish CC, Durstewitz D, Chandler LJ, Seamans JK. Successful choice behavior is associated with distinct and coherent network states in anterior cingulate cortex. Proc Natl Acad Sci. 2008;105(33):11963–11968. doi:10.1073/pnas.0804045105.

37. Durstewitz D. Implications of synaptic biophysics for recurrent network dynamics and active memory. Neural Networks. 2009;22(8):1189–1200.

38. Izhikevich EM. Dynamical systems in neuroscience. MIT press; 2007.

39. Durstewitz D. Self-Organizing Neural Integrator Predicts Interval Times through Climbing Activity. J Neurosci. 2003;23(12):5342–5353.

40. Markram H, Toledo-Rodriguez M, Wang Y, Gupta A, Silberberg G, Wu C. Interneurons of the neocortical inhibitory system. Nat Rev Neurosci. 2004;5(10):793–807. doi:10.1038/nrn1519.

41. Kamigaki T, Dan Y. Delay activity of specific prefrontal interneuron subtypes modulates memory-guided behavior. Nature neuroscience. 2017;20(6):854.

42. Kriener B, Enger H, Tetzlaff T, Plesser HE, Gewaltig MO, Einevoll GT. Dynamics of self-sustained asynchronous-irregular activity in random networks of spiking neurons with strong synapses. Frontiers in computational neuroscience. 2014;8:136.

43. Lundqvist M, Compte A, Lansner A. Bistable, irregular firing and population oscillations in a modular attractor memory network. PLoS Computational Biology. 2010;6(6):e1000803.

44. Van Vreeswijk C, Sompolinsky H. Chaos in neuronal networks with balanced excitatory and inhibitory activity. Science. 1996;274(5293):1724–1726.

45. Brecht M, Sakmann B. Dynamic representation of whisker deflection by synaptic potentials in spiny stellate and pyramidal cells in the barrels and septa of layer 4 rat somatosensory cortex. J Physiol. 2002;543:49–70.

46. Shoham S, O’Connor DH, Segev R. How silent is the brain: is there a “dark matter” problem in neuroscience? J Comp Physiol [A]. 2006;192:777–784.

47. Barth AL, Poulet JFA. Experimental evidence for sparse firing in the neocortex. Trends Neurosci. 2012;35(6):345–355.

48. Durstewitz D, Seamans JK, Sejnowski TJ. Dopamine-Mediated Stabilization of Delay-Period Activity in a Network Model of Prefrontal Cortex. J Neurophysiol. 2000;83:1733–1750.

49. Hass J, Durstewitz D. Models of dopaminergic modulation. Scholarpedia. 2011;6(8):4215.

50. Compte A, Constantinidis C, Tegnér J, Raghavachari S, Chafee MV, Goldman-Rakic PS, et al. Temporally irregular mnemonic persistent activity in prefrontal neurons of monkeys during a delayed response task. Journal of neurophysiology. 2003;90(5):3441–3454.

51. Khan AG, Poort J, Chadwick A, Blot A, Sahani M, Mrsic-Flogel TD, et al. Distinct learning-induced changes in stimulus selectivity and interactions of GABAergic interneuron classes in visual cortex. Nature neuroscience. 2018; p. 1.

52. Marder E. Variability, compensation, and modulation in neurons and circuits. Proceedings of the National Academy of Sciences. 2011;108(Supplement 3):15542–15548.

53. Vitureira N, Letellier M, Goda Y. Homeostatic synaptic plasticity: from single synapses to neural circuits. Current opinion in neurobiology. 2012;22(3):516–521.

54. Börgers C, Epstein S, Kopell NJ. Background gamma rhythmicity and attention in cortical local circuits: a computational study. Proceedings of the National Academy of Sciences of the United States of America. 2005;102(19):7002–7007.

55. Lundqvist M, Rose J, Herman P, Brincat SL, Buschman TJ, Miller EK. Gamma and beta bursts underlie working memory. Neuron. 2016;90(1):152–164.

56. Russo E, Durstewitz D. Cell assemblies at multiple time scales with arbitrary lag constellations. eLife. 2017;6:e19428.

57. Miller P, Brody CD, Romo R, Wang XJ. A recurrent network model of somatosensory parametric working memory in the prefrontal cortex. Cerebral Cortex. 2003;13(11):1208–1218.

58. Rabinovich M, Volkovskii A, Lecanda P, Huerta R, Abarbanel H, Laurent G. Dynamical encoding by networks of competing neuron groups: winnerless competition. Physical review letters. 2001;87(6):068102.

59. Lundqvist M, Herman P, Lansner A. Theta and gamma power increases and alpha/beta power decreases with memory load in an attractor network model. Journal of cognitive neuroscience. 2011;23(10):3008–3020.

60. Siegel M, Warden MR, Miller EK. Phase-dependent neuronal coding of objects in short-term memory. Proceedings of the National Academy of Sciences. 2009;106(50):21341–21346.

61. Buschman TJ, Denovellis EL, Diogo C, Bullock D, Miller EK. Synchronous oscillatory neural ensembles for rules in the prefrontal cortex. Neuron. 2012;76(4):838–846.

62. Kornblith S, Buschman TJ, Miller EK. Stimulus load and oscillatory activity in higher cortex. Cerebral Cortex. 2015; p. bhv182.

63. Brette R, Gerstner W. Adaptive exponential integrate-and-fire model as an effective description of neuronal activity. J Neurophysiol. 2005;94:3637–3642.

64. Gerstner W, Kistler WM. Spiking Neuron Models: Single Neurons, Populations, Plasticity. Cambridge University Press; 2002.

65. Krimer LS, Goldman-Rakic PS. Prefrontal Microcircuits: Membrane Properties and Excitatory Input of Local, Medium, and Wide Arbor Interneurons. J Neurosci. 2001;21(11):3788–3796.

66. Song S, Sjöoströom PJ, Reigl M, Nelson S, Chklovskii DB. Highly Nonrandom Features of Synaptic Connectivity in Local Cortical Circuits. PLoS Biol. 2005;3(3):e68. doi:10.1371/journal.pbio.0030068.

67. Maass W, Markram H. Synapses as dynamic memory buffers. Neural Networks. 2002;15(2):155–161. doi:10.1016/S0893-6080(01)00144-7.

