## Supplemental Materials for "Constraints on Persistent Activity in a Biologically Detailed Network Model of the Prefrontal Cortex with Heterogeneities"

**Supporting figure S1**

**Note:** The figure is taken from Hass, Hass J, Hertäg L, Durstewitz D (2016) A Detailed Data-Driven Network Model of Prefrontal Cortex Reproduces Key Features of In Vivo Activity. PLoS Comput Biol 12(5): e1004930. doi:10.1371/journal.pcbi.1004930.

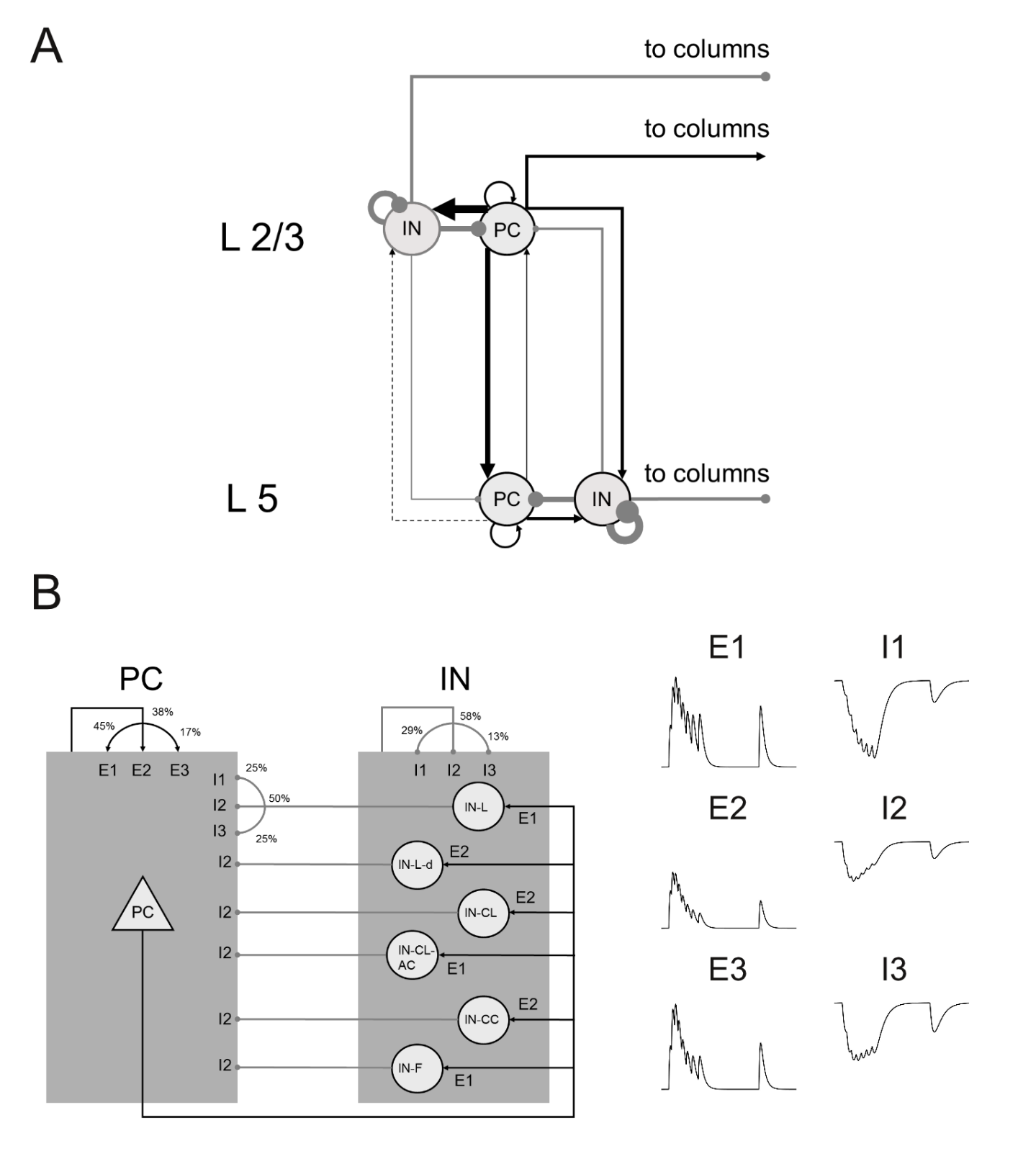

**Supporting tables S1-S4**

**Note:** All tables are taken from Hass, Hass J, Hertäg L, Durstewitz D (2016) A Detailed Data-Driven Network Model of Prefrontal Cortex Reproduces Key Features of In Vivo Activity. PLoS Comput Biol 12(5): e1004930. doi:10.1371/journal.pcbi.1004930. DOIs for the individual tables are provided.

**Supporting table S1. Neuron parameters**

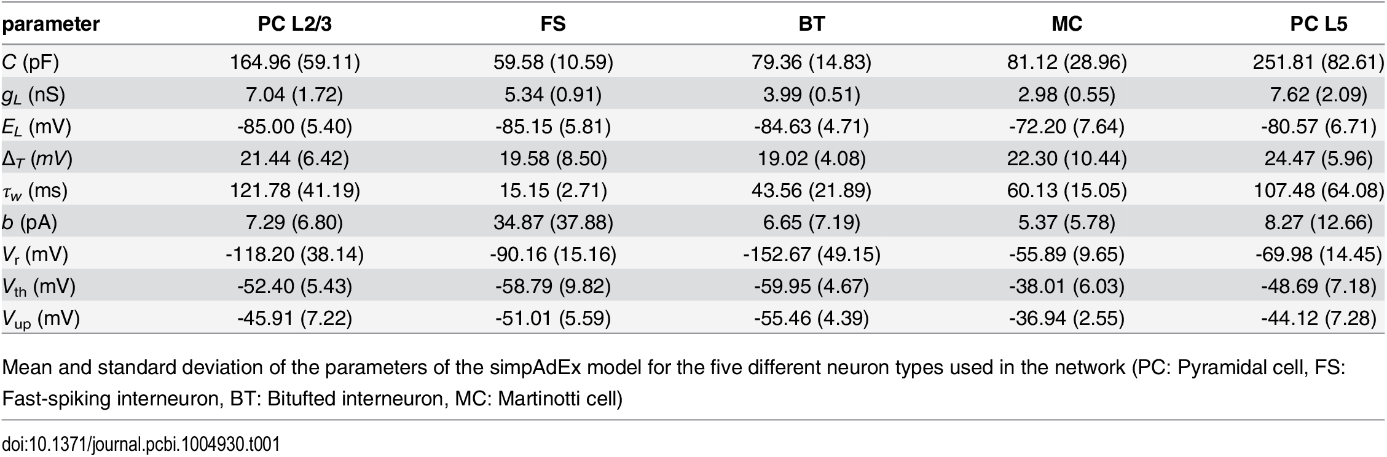

**Supporting table S2. Cell numbers**

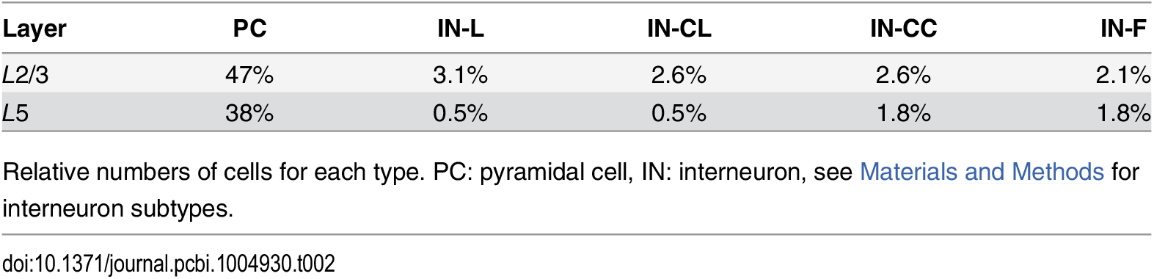

**Supporting table S3. Synaptic parameters**

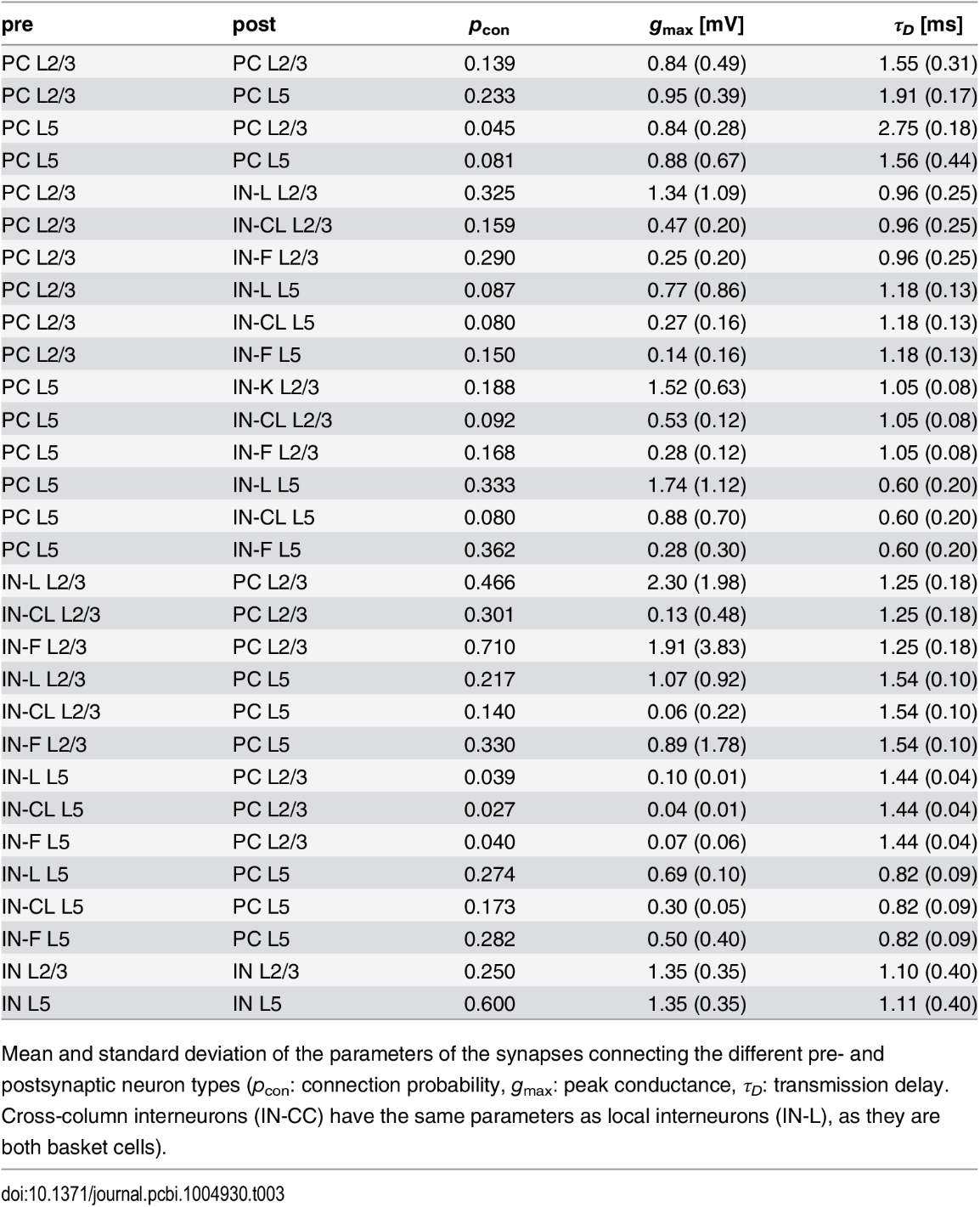

**Supporting table S4. Short-term synaptic plasticity parameters**

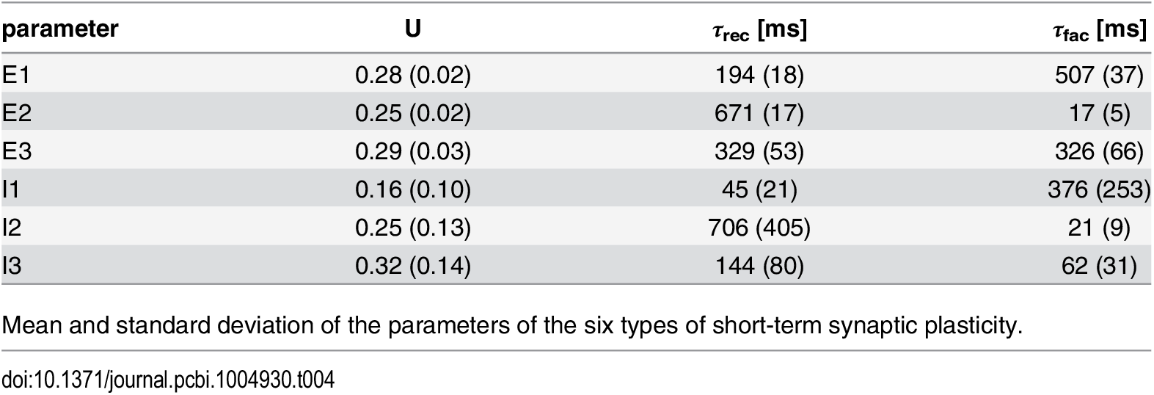
